# The RhoGEF Cysts couples apical polarity proteins to Rho and myosin activity at adherens junctions

**DOI:** 10.1101/617134

**Authors:** Jordan T. Silver, Frederik Wirtz-Peitz, Sérgio Simões, Milena Pellikka, Dong Yan, Richard Binari, Takashi Nishimura, Yan Li, Tony J. C. Harris, Norbert Perrimon, Ulrich Tepass

## Abstract

The spatio-temporal regulation of small Rho GTPases is crucial for the dynamic stability of epithelial tissues. However, how RhoGTPase activity is controlled during development remains largely unknown. To explore the regulation of Rho GTPases in vivo we analyzed the Rho GTPase guanine nucleotide exchange factor (RhoGEF) Cysts, the Drosophila orthologue of mammalian p114RhoGEF, GEF-H1, p190RhoGEF, and AKAP-13. Loss of Cysts causes a phenotype that closely resemble the mutant phenotype of the apical polarity regulator Crumbs. This phenotype can be suppressed by the loss of basolateral polarity proteins suggesting that Cysts in an integral component of the apical polarity protein network. Cysts activates Rho at adherens junctions to promote junctional enrichment of myosin II, which requires the RhoGEF domain and the coiled-coil domain containing C-terminal region of Cysts. Cysts recruitment to the apico-lateral cortex depends on Crumbs and Bazooka/Par3 and requires multiple domains within Cysts including the C-terminal region. Together, our findings indicate that Cysts links apical polarity proteins to Rho1 and myosin activation at adherens junctions to support junctional and epithelial integrity in the Drosophila ectoderm.

## Introduction

Epithelial cells show pronounced apical-basal polarity and form an apical junctional complex that encircles individual cells and tightly links neighboring cells into a sheet-like tissue. Factors that regulate apical-basal polarity, and in particular apical polarity proteins such as Crumbs (Crb) or atypical protein kinase C (aPKC) play a pivotal role in the formation of the junctional belt so that the loss of either core junctional proteins such as E-cadherin or apical polarity proteins causes a similar loss of epithelial integrity (Tepass, 2012). How the function of apical polarity proteins support a circumferential junctional belt is not well-understood. One key mechanisms that promotes junctional stability is the activity of cytoplasmic myosin II that is activated at apical junctions through the Rho-Rock pathway (Mack and Georgiou, 2014; Lecuit and Yap, 2015). Here, we ask how apical polarity proteins help to confine the activity of Rho1 to the apical junctional region to promote the formation of a junctional belt.

Rho GTPases are molecular switches that cycle between active (GTP-bound) and inactive (GDP-bound) states. Rho GTPases are utilized over and over again to regulate diverse molecular processes within cells including cytoskeletal dynamics, cell polarity, cell adhesion, and vesicle trafficking (Jaffe and Hall, 2005; Hall, 2012; Ridley, 2012; Ratheesh and Priya, 2013; Mack and Georgiou, 2014). For example, epithelial polarity in the Drosophila embryo requires three members of the Rho protein family: Cdc42 activates the apical Par protein complex (Par6/aPKC) to promote apical polarity (Hutterer et al., 2004; Harris and Tepass, 2008; Harris and Tepass, 2010), Rac1 acts together with phosphoinositide 3-kinase (PI3K) as a basolateral polarity protein (Chartier et al., 2011), and Rho1 (RhoA in mammals) supports integrity of apical adherens junctions (AJs) by activating myosin II through the Rho Kinase (Rok) pathway (Magie et al., 1999; Fox et al., 2005; Matsuoka and Yashiro, 2014; Lecuit and Yap, 2015).

Rho GTPase-specific guanine nucleotide exchange factors (RhoGEFs) and Rho GTPase-activating proteins (RhoGAPs) promote activation and deactivation of Rho GTPases respectively (McCormack et al. 2013; Cook et al., 2014). The Drosophila genome encodes 26 RhoGEFs and 22 RhoGAPs that presumably regulate seven Rho GTPases and a wide variety of downstream effectors, many of which modulate cytoskeletal remodelling (Aspenström, 1999; Greenberg and Hatini, 2011; Hall, 2012; Cook et al., 2014). Rho1, Cdc42, and Rac (with three fly paralogs Rac1, Rac2, and Mtl) are maternally provided to the embryo (Magie et al., 1999, Genova et al., 2000, Hakeda-Suzuki et al., 2002). These Rho GTPases play multiple essential roles in embryogenesis through their contributions to epithelial polarity, cell movements such as mesoderm invagination, germband extension, dorsal closure, and wound repair among other processes (Harden, 2002; Mack and Georgiou, 2014; Verboon and Parkhurst, 2015). However, how RhoGTPases are regulated in development through the spatial and temporal engagement of GEFs and GAPs still remains largely unexplored.

Here, we characterize a Drosophila RhoGEF Cysts (*cyst*; *CG10188*). Cyst is the single orthologue of a group of four mammalian paralog GEFs characterized by the presence of a DH-PH domain (RhoGEF or Dbl homology-pleckstrin homology): p114RhoGEF, p190RhoGEF, AKAP-13, and GEF-H1 (McCormack et al., 2013; Cook et al. 2014; Ngok et al., 2014). Tissue culture studies have shown that p114RhoGEF (ARHGEF18 in humans) links the epithelial polarity machinery with the actomyosin cytoskeleton. p114RhoGEF interacts with proteins of the apical Par and Crb polarity complexes and activates RhoA in support of actomyosin organization, apical constriction, and junction assembly (Nakajima and Tanoue, 2010; 2011; 2012; Terry et al., 2011; Acharya et al., 2018). Whether p190RhoGEF, AKAP-13, or GEF-H1 play a role in epithelial polarity is currently unknown. Our work shows that Cyst is a key RhoGEF in the Drosophila embryo that targets Rho1 and consequently myosin II at AJs to support epithelial apical-basal polarity. In addition to an essential RhoGEF domain, we show that the C-terminal region of Cyst plays an important role in recruiting Cyst to the apico-lateral cortex. Our data suggest that Patj, Bazooka (Baz, the Drosophila homolog of Par3), and Crb control the polarized distribution of Cyst. We propose that Cyst is a crucial component of the apical polarity protein network that couples apical polarity to the junctional Rho1-myosin pathway to support AJ stability and epithelial integrity.

## Results

### Cyst is an apical epithelial polarity protein required for AJ stability during gastrulation

Maternal knockdown of *CG10188* caused an embryonic lethal phenotype characterized by the formation of many small epithelial vesicles or cysts instead of a large epithelial sheet. We named *CG10188* therefore *cysts* (*cyst*) (Figure 1C). These *cyst* RNAi embryos also displayed shields of dorsal cuticle. *Cyst* RNAi embryos were rescued to adulthood by a *Cyst^R^* transgene, containing a *cyst* genomic sequence that has been mutagenized to render it immune to *cyst* RNAi without altering the encoded protein sequence (Figure 1A). We also noted that overexpression of Cyst led to embryonic lethality with embryos displaying severe defects in head morphogenesis (Figure 1F). These results identify Cyst as an essential factor for maintaining epithelial integrity in the Drosophila embryo.

**Figure 1.**
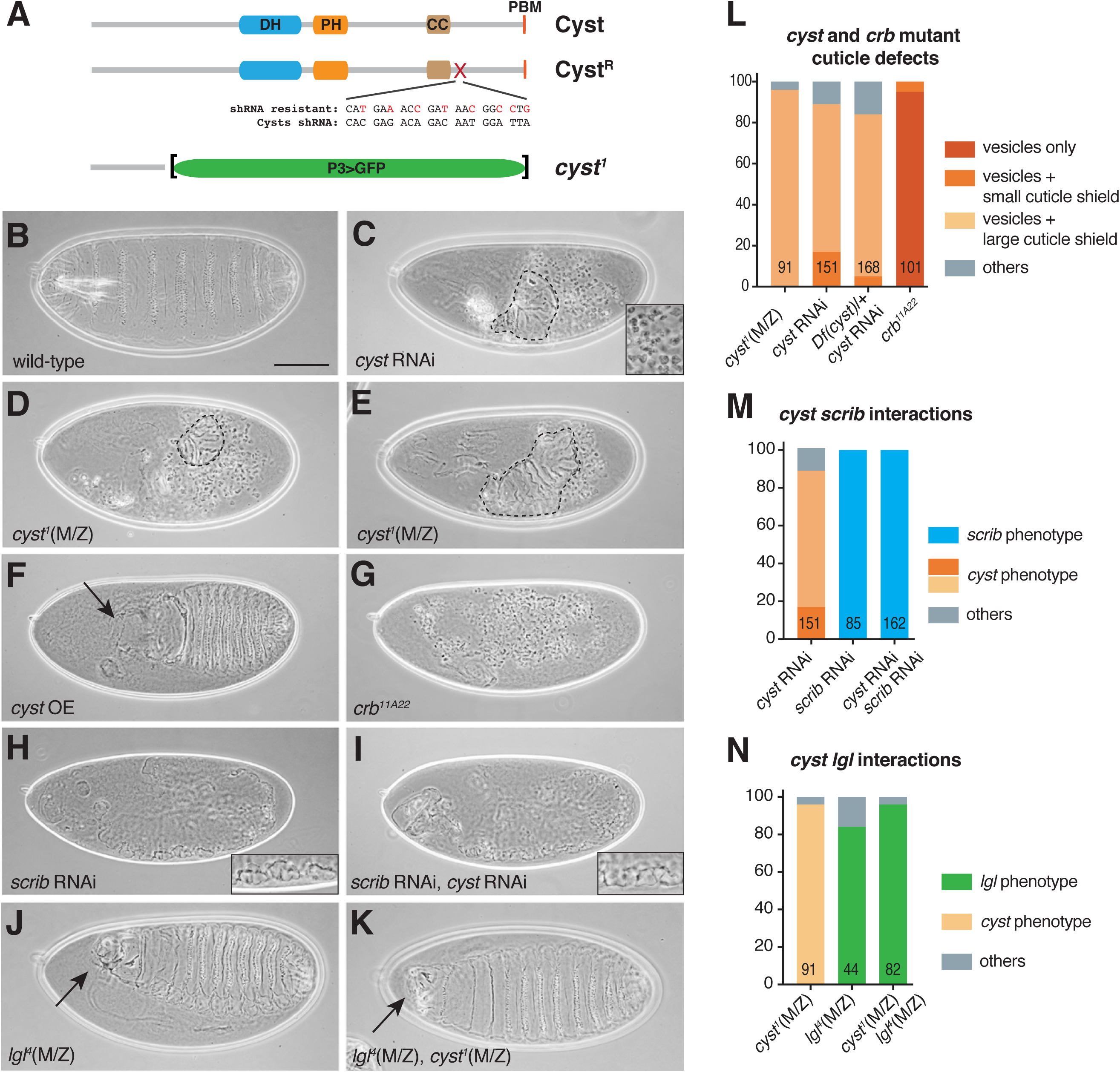
Cyst maintains epithelial polarity. **(A)** Schematic of Cyst and Cyst^R^ (shRNA resistant) and *cyst^1^*. Cyst^R^ is encoded by a *cyst* genomic sequence that has been mutagenized to render it immune to *cyst* shRNA. Cyst is expressed under the endogenous promoter. DH – Dbl homology; PH – pleckstrin homology; CC – coiled coil; PBM – PDZ-binding motif. *cyst^1^* is a genomic deletion where *cyst* sequence has been replaced by GFP driven by the P3 promoter. **(B-K)** Cuticles of embryos of the indicated genotypes. *cyst* RNAi (C,I) and *scrib* RNAi (H,I) embryos derived from mothers expressing shRNAs with mat-Gal4. *cyst^1^*(M/Z) *lgl^4^*(M/Z) are *cyst^1^* mutant (D,E,K) or *lgl^4^* mutant (J,K) germline clones. *cyst* OE (overexpression), mat-Gal4 UAS-*cyst* embryo (F). *crb^11A22^*, *crb* null mutant embryo (G). Inset in (C) shows field of cuticle vesicles/cysts. Inset in (H) and (I) show the bubbly cuticle defects typical for *scrib* depleted embryos. Dashed lines encircle cuticle shields (C-E). Arrows point to head defects. Scale bar, 100 μm (B-K). **(L-N)** Quantification of cuticle defects of indicated genotypes.

We generated a *cyst* deletion mutation (*cyst^1^*) with CRISPR/Cas9 technology that removes most of the *cyst* coding sequence including the DH (RhoGEF) and PH domains, and the entire C-terminal part of the protein (Figure 1A). *cyst^1^* mutant animals are not embryonic lethal. In contrast, embryos derived from *cyst^1^* mutant germ line clones (*cyst*^M/Z^ embryos) did show a prominent embryonic phenotype similar to *cyst* RNAi embryos, characterized by the presence of a large number of epithelial vesicles and shields of cuticle of various sizes (Figure 1 D,E). The *cyst* phenotype is reminiscent of the *crb* mutant phenotype (Figure 1G) (Tepass et al., 1990; Tepass and Knust, 1990; 1993) or the phenotype caused by depletion of other apical polarity proteins or AJ proteins such as DE-cadherin (DEcad) (Tepass et al., 1996, Uemura et al., 1996). Whereas most *crb* mutant embryos only showed cuticle vesicles, all *cyst^M/Z^* or *cyst* RNAi embryos showed cuticle shields in addition to vesicles suggesting that the *cyst* phenotype is qualitatively similar but somewhat weaker than the *crb* phenotype.

A *crb*-like phenotype has been documented for only a small number of genes that encode apical polarity proteins such as *crb* and *stardust* (*sdt*), or genes encoding AJ proteins such as *bazooka* (*baz*), *shotgun* (encoding DEcad) or *armadillo* (*arm*; Drosophila β-catenin) (Tepass et al., 1990, Tepass and Knust, 1990; 1993; Tepass et al., 1996; Cox et al., 1996; Tanentzapf and Tepass, 2003; Bilder et al., 2003). In contrast, genes encoding basolateral polarity proteins such as *scribble* (*scrib*) or *yurt* (*yrt*) have different mutant phenotypes and do not display epithelial cysts with inward facing apical lumina (Bilder et al., 2000; Tanentzapf and Tepass, 2003; Bilder et al., 2003; Laprise et al., 2006; 2009). The cuticle defects observed in *cyst* compromised embryos therefore strongly suggest that Cyst is a new component of the apical polarity machinery. The development of a *crb*-like phenotype was described in detail and entails the loss of polarity and AJ fragmentation during gastrulation (stages 8-11). Subsequently, these embryos show enhanced programmed cell death elicited by activation of the JNK signaling pathway (Kolahgar et al., 2011). Epithelial cells that survive form cysts which show normal polarity and secret a luminal cuticle (Tanentzapf and Tepass, 2003). Suppression of cell death amplifies the number of cysts suggesting that even cells that die do not lack the ability to form polarized epithelial units (Tanentzapf and Tepass, 2003). To determine whether a similar sequence of events can be found in *cyst* compromised embryos we examined the development of *cyst* embryos with live imaging monitoring the AJ marker DEcad::GFP (Figure 2), and by assessing the distribution of apical (Crb, aPKC), junctional (Arm), and basolateral (Yrt) polarity markers (Figure 3 and S1). These observations showed progressive AJ fragmentation during gastrulation until many cells in particular in the ventral ectoderm had lost AJs (Figure 2 A,B). Other cells showed focal concentrations of DEcad suggesting the beginning of cyst formation (Figure 2C). Cyst formation is first seen at the end of gastrulation in stage 11 embryos and can be followed throughout the rest of development (Figure 3 and S1). Cysts showed normal polarized distribution of Crb, aPKC, Arm and Yrt (Figure 3 and S1). Collectively, these findings indicate that Cyst function is closely related to apical and/or junctional epithelial polarity regulators.

**Figure 2.**
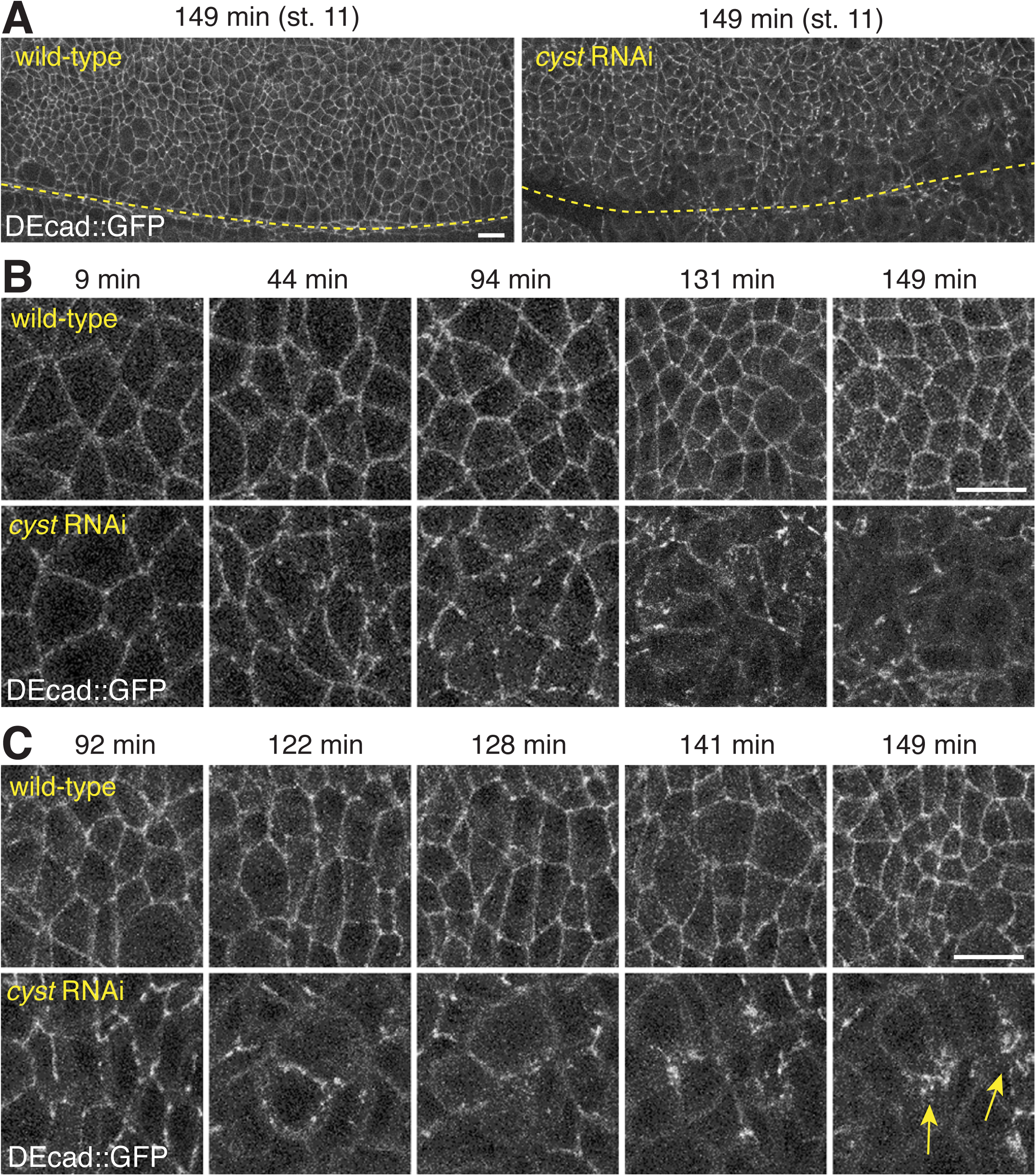
Cyst is required for AJ integrity. Z-projections (9.5 μm) taken from live wild type or *cyst* RNAi embryos expressing endoDEcad::GFP (see Movies S1 and S2). Times indicate minutes after onset of germband extension. Scale bars, 10 μm. **(A)** Ventral view of the ectoderm at stage 11 showing a loss of AJs in cells adjacent to the ventral midline (dashed line) in *cyst* RNAi embryo. **(B)** Ventral ectoderm cells at the indicated time points showing the increasing fragmentation and loss of AJs in *cyst* RNAi embryo. **(C)** Ventral ectoderm cells at the indicated time points showing clustering of AJ material in *cyst* RNAi embryo (arrows).

**Figure 3.**
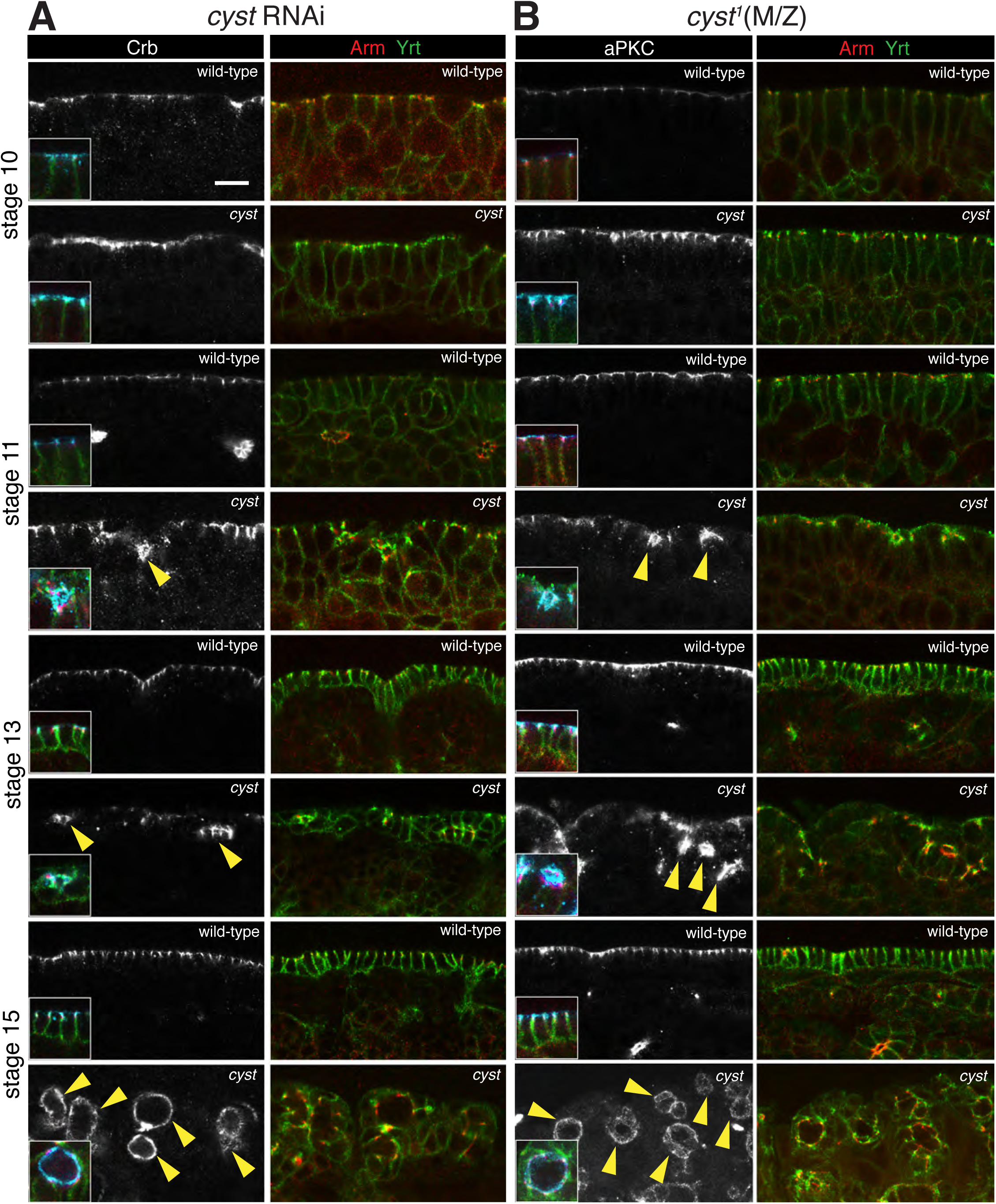
Cysts forming in *cyst* compromised embryos maintain epithelial polarity. **(A,B)** Staining of *cyst* RNAi (A) and *cyst^1^*(M/Z) (B) embryos compared to wild-type controls for the apical markers Crb and aPKC, the junctional marker Arm, and the basolateral marker Yrt. All panels show side views of the ectoderm or epidermis at the indicated stages. Cysts in *cyst* compromised embryos are evident in stage 11, 13 and 15 embryos (arrowheads) but not at stage 10. Markers show normal subcellular distributions in cysts with apical markers facing the lumen where cuticle will be secreted (see Figure 1 C-E). Insets show triple labeled cells with Crb or aPKC shown in blue. Scale bar, 10 μm.

A key feature of the machinery that regulates epithelial polarity is the negative feedback between apical and basolateral polarity proteins (Tanentzapf and Tepass, 2003; Bilder et al, 2003; Benton and St Johnston, 2003; Laprise et al., 2009; Chartier et al., 2011; Fletcher et al., 2012; Gamblin et al., 2014). This mutual antagonism can be revealed through double mutant analysis. For example, double mutants of *crb* and the basolateral polarity gene *scrib* or *sdt* and the basolateral polarity gene *lethal giant larvae* (*lgl*) show a striking suppression of the *crb* or *sdt* mutant defects and display a *scrib* or *lgl* mutant phenotype, respectively (Tanentzapf and Tepass, 2003; Bilder et al., 2003). To reveal whether *cyst* behaves like *crb* or *sdt* in these tests, we generated embryos compromised for *cyst* and *scrib* or *cyst* and *lgl* (Figure 1H-K,M,N). Maternal depletion of *scrib* with RNAi caused a strong loss-of-function phenotype characterized by epidermal cell clusters surrounded by cuticle (Figure 1H) (Tanentzapf and Tepass, 2003; Bilder et al, 2003), whereas *lgl^4^*(M/Z) embryos displayed a weaker phenotype with defects in head morphogenesis. *scrib* RNAi fully suppressed the *cyst* RNAi phenotype and *lgl^4^*(M/Z) fully suppressed the *cyst^1^*(M/Z) phenotype with double mutant showing phenotypes indistinguishable from *scrib* RNAi or *lgl^4^*(M/Z) alone (Figure 1I,M,K,N). Taken together, our analysis suggests that Cyst is a new key component of the apical polarity machinery that acts during gastrulation to maintain junctional and epithelial integrity in the fly embryos.

### Cyst regulates planar epithelial organization in the early embryo

To further assess the role of Cyst in the early embryo, we analyzed the lateral ectoderm at the onset of germband extension, when circumferential AJs form. At this stage, DEcad, Baz and other AJ proteins become enriched at the apico-lateral boundary but also acquire a planar polarized distribution to dorsal and ventral edges of the apico-lateral domain (Zallen and Wieschaus, 2004). Disrupted planar polarization, with Baz hyper-polarizing as prominent, single foci along cell edges have been observed when actin is reduced or in embryos with abnormal activity of polarity proteins including aPKC, Par1, and Crb (Harris and Peifer, 2007; Jiang et al., 2015; Vichas et al., 2015). This tissue thus provides a context to examine whether Cyst cooperates with polarity proteins in the initial formation of a normal circumferential AJ belt.

We found that *cyst* depletion by maternal expression of shRNA alone had minimal effects on Baz distribution (data not shown). However, further depletion of *cyst* by maternal heterozygosity for a deletion comprising *cyst* (*Df(cyst)* = *Df(2L)BSC301*), produced Baz hyper-polarization in contrast to control (mCherry shRNA) or *Df(cyst)*/+ alone (Figure 4A-B, not shown). To test if Baz hyper-polarization was subject to regulation by aPKC, we analyzed *cyst* RNAi*; Df(cyst)*/+ embryos derived from mothers heterozygous for a null aPKC allele, and discovered even greater Baz hyper-polarization, whereas *Df(cyst)*/+, *apkc*/+ embryos were normal (Figure 4A-B). As compromising the Cdc42-binding region of Par6 disrupts the cortical localization of the Par6/aPKC complex (Hutterer et al., 2004), we investigated whether the mis-regulation of Baz with *cyst* depletion could be explained by a loss of aPKC from the apical domain. However, *cyst* loss-of-function embryos displayed no detectable effects on aPKC localization when the blastoderm is fully formed and before the germband starts to extend (Figure 4C), or on aPKC levels around the margins of the apical domain (Figure 4C-D). As an alternative possibility, we examined if disruption of F-actin could explain the observed hyperpolarization of Baz. Treatment of embryos with cytochalasin D resulted in Baz hyper-polarization (Figure 4E), with no apparent effect on aPKC levels at the apico-lateral membrane (Figure 4F), mimicking the effects of *cyst* loss-of-function. These results indicate that Cyst contributes to the initial formation of a normal circumferential AJ belt. Cyst appears to act by regulating the actin cytoskeleton, either downstream or in parallel to an aPKC-dependent mechanism.

**Figure 4.**
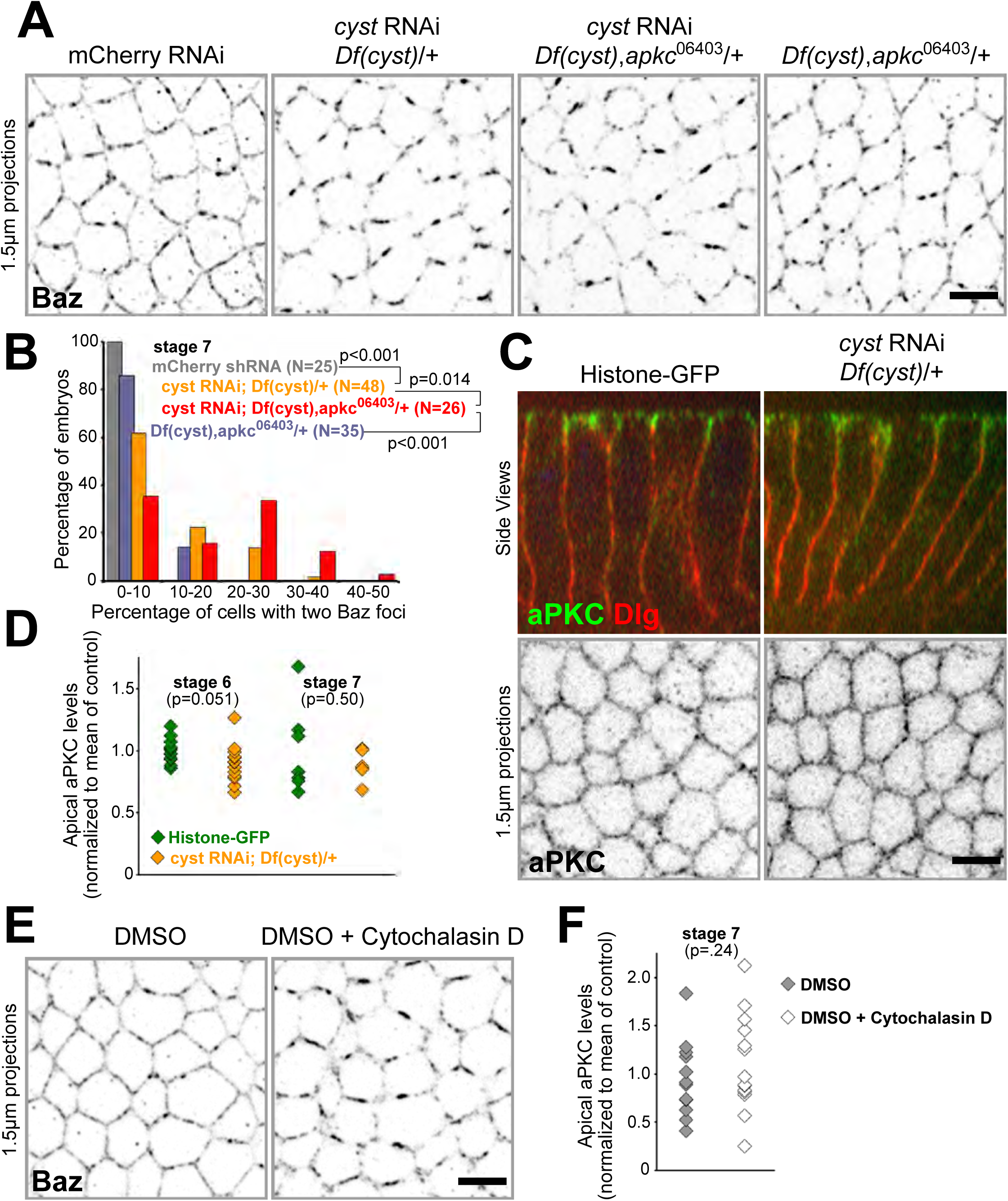
Cyst contributes to the initial formation of an AJ belt. **(A)** Immunostaining of Baz in the apico-lateral domain of the lateral ectoderm at the onset of germband extension. Reduction of *cyst* expression through the combination of maternal shRNA (using MTD-GAL4) and maternal heterozygosity for *Df(cyst)* resulted in Baz planar hyper-polarization into foci. Additional incorporation of maternal heterozygosity for a null *apkc* allele enhanced the hyper-polarization of Baz. A control shRNA, or the combined heterozygosity of *Df(cyst)* and *apkc*, had no apparent effects. Scale bar, 5 μm. **(B)** Quantification of the data shown in (A). N-values represent embryo numbers. **(C)** Side views (top) and apical domain surface projections (bottom) showing indistinguishable apical polarization and apical levels of aPKC with Cyst depletion versus internal controls at the onset of germband extension. Dlg labels the lateral domain. Scale bar, 5 μm. **(D)** Quantification of the data shown in (C). Each point is a quantification of one embryo. **(E)** Cytochalasin D produces Baz hyper-polarization at the onset of germband extension gastrulation, in contrast to carrier DMSO control. Scale bar, 5 μm. **(F)** Cytochalasin D has no apparent effect on apical aPKC levels at the onset of germband extension. Each point is a quantification of one embryo.

### Cyst localizes to the apico-lateral cortex

The impact of Cyst on AJ formation suggests that it may localize to the apico-lateral cortex to exert its function. To ask whether Cyst is found at AJs we examined the distribution of an N-terminally GFP-tagged isoform of Cyst (GFP::Cyst) expressed under the *cyst* endogenous promoter in live embryos. GFP::Cyst was indeed enriched at the apico-lateral cortex (Figure 5A,B). We found that GFP::Cyst concentration increases at the cortex from stage 8 to mid-embryogenesis (stage 14) and decreases at later stages. These findings suggest that Cyst is enriched at AJs or their immediate vicinity from the onset of germband extension (stage 6/7) and throughout organogenesis.

**Figure 5.**
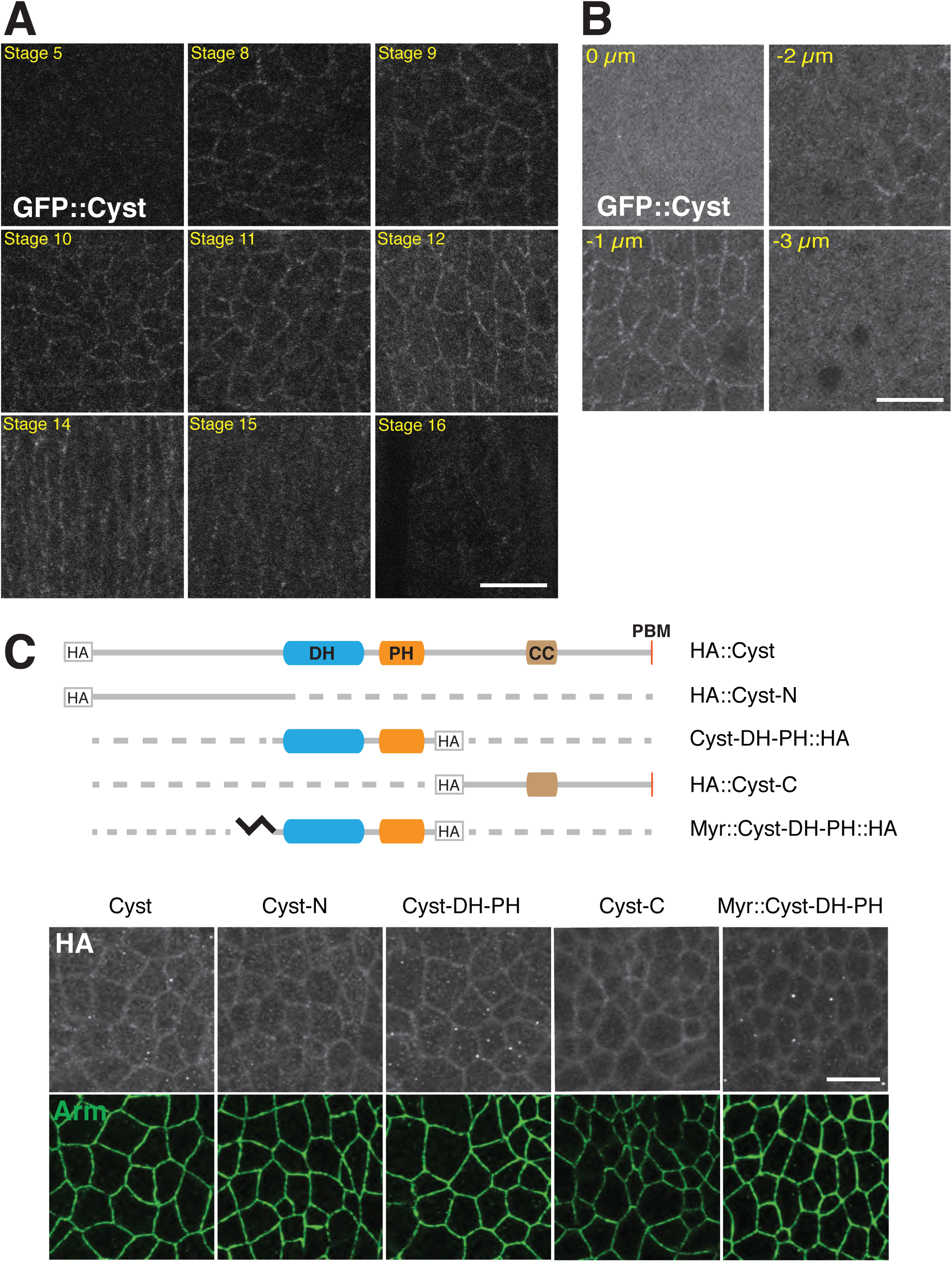
Cyst localizes to the apico-lateral cortex of epithelial cells. **(A)** Z-projections (1 μm deep) of the ventral ectoderm were assembled from apical planes taken from live *cyst* RNAi embryos rescued by GFP::Cyst. GFP::Cyst is enriched at the apico-lateral cortex. GFP::Cyst levels are low before stage 8, increase from stage 8 to 12, and decrease from stage 15 and onwards. Scale bar, 10 μm. **(B)** Serial z-projections (1 μm deep) of the ventral ectoderm from a wild-type stage 11 embryo expressing GFP::Cyst. 0 μm shows the vitelline membrane. GFP::Cyst is enriched at the cortex 1 μm below (−1 μm) the vitelline membrane. Scale bar, 10 μm. **(C)** Schematic of Cyst overexpression constructs. Constructs are HA-tagged and expressed under the *UASp* promoter. Black zigzag – myristoylation; dashed line – deleted regions. Immunostainings (anti-HA and anti-Armadillo; Arm) of stage 11 embryos expressing the indicated constructs. 1 μm deep z-projections at the level of the AJs of the ventral ectoderm are shown. All constructs are cortically enriched and overlap with Arm. Scale bar, 10 μm.

### Cyst contains multiple regions that can facilitate junctional localization

We conducted a structure-function analysis of Cyst to better understand the mechanism underlying Cyst polarized distribution. Similar to GFP::Cyst, we found that a UAS-controlled maternally loaded N-terminal HA-tagged version of Cyst (HA::Cyst) was enriched at the level of AJs as shown for stage 11 embryos (Figure 5C). Interestingly, N-, DH-PH-, and C-fragments of Cyst (Cyst-N, Cyst-DH-PH, and Cyst-C) were also enriched at the level of AJs. Myristoylation of the DH-PH domain (Myr::Cyst-DH-PH) did not further enhance cortical association (Figure 5C). Together, these results suggest that Cyst has multiple membrane targeting sequences.

To further test the hypothesis that multiple regions of Cyst can facilitate its association with the cell cortex, we examined the activity and distribution of shRNA-resistant GFP-tagged Cyst genomic structure-function constructs (Figure 6A). Constructs were expressed both in a *cyst* shRNA background and in wild-type embryos. Notably, all constructs were detected at the apico-lateral cortex when expressed in *cyst* shRNA embryos although the distribution of Cyst^R^ΔDH-PH::GFP, Cyst^R^ΔC::GFP and GFP::Cyst^R^ΔPBM (lacking DH-PH domain responsible for the GEF activity, the C-terminal region, or putative C-terminal PDZ-binding motif at 1306-EIYF-1309) was more punctate than GFP::Cyst^R^ or GFP::Cyst^R^ΔN (Figure 6D). Moreover, we found that GFP::Cyst^R^, GFP::Cyst^R^ΔN, and Cyst^R^ΔDH-PH::GFP were enriched at the apico-lateral cortex in the presence of endogenous Cyst protein (Figure 6D). In contrast, Cyst^R^ΔC::GFP and GFP::Cyst^R^ΔPBM were not found at the cortex in the presence of endogenous Cyst. This finding suggests that endogenous Cyst out-competes Cyst^R^ΔC::GFP and GFP::Cyst^R^ΔPBM. Together, our analysis of genomic and overexpression constructs reveals that Cyst contains multiple regions that facilitate its apically polarized distribution. In particular, the Cyst C-terminal region harbours strong localization sequences.

**Figure 6.**
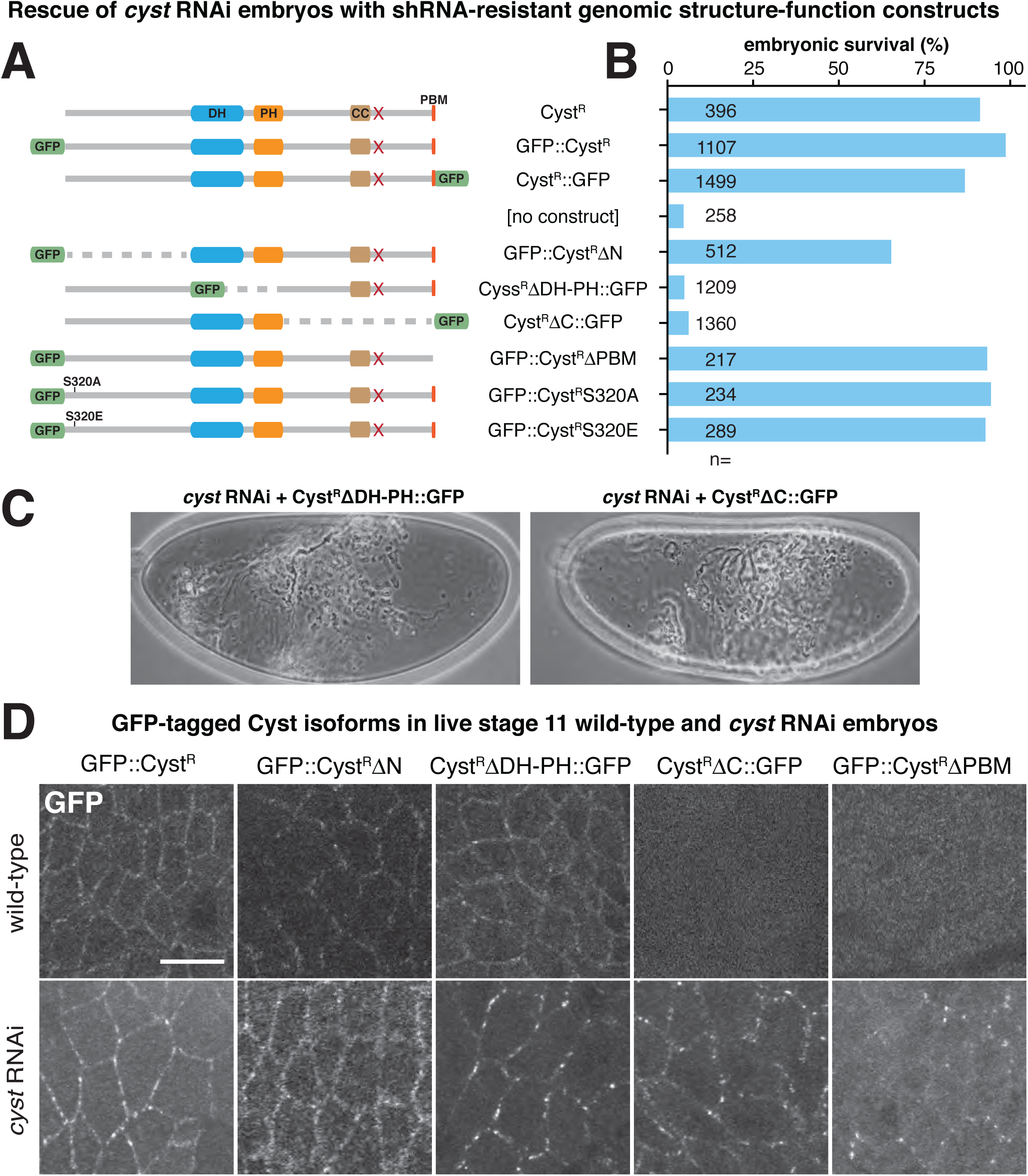
Cyst contains essential RhoGEF and C-terminal regions. **(A)** Schematic of Cyst GFP-tagged structure-function constructs. Constructs are generated from the *cyst* genomic region, expressed under the endogenous promoter, and are immune to *cyst* shRNA (red cross; see figure 1A)). GFP replaces deleted regions (dashed lines). **(B)** Quantification of embryonic rescue activity of Cyst isoforms expressed in a *cyst* RNAi background. Cyst^R^ΔN partially rescues *cyst* RNAi embryos (68% embryonic viable; *p<0.001 compared to Cyst^R^ control). **(C)**Embryo cuticles of the indicated genotypes showing that Cyst^R^ΔDH-PH and Cyst^R^ΔC fail to rescue the *cyst* RNAi phenotype. **(D)** Z-projections (1 μm deep) ventral ectoderm taken from live stage 11 wild-type or *cyst* RNAi embryos expressing the indicated constructs. All Cyst isoforms are cortically enriched in a *cyst* RNAi background. In wild-type embryos, Cyst^R^ΔC::GFP and Cyst^R^ΔPBM were not detected at the cortex. Scale bar, 10 μm.

### The RhoGEF domain and the C-terminal region are essential for Cyst function

shRNA-resistant genomic structure-function constructs were also tested for their ability to rescue *cyst* RNAi embryos (Figure 6A,B). As expected, a construct lacking the DH-PH domain of Cyst (Cyst^R^ΔDH-PH::GFP) did not rescue *cyst* RNAi embryos in contrast to control constructs (Cyst^R^, Cyst^R^::GFP, GFP::Cyst^R^; Figure 6A,B), indicating that the RhoGEF domain of Cyst is essential for epithelial integrity. In line with our finding that Cyst contains C-terminal sequences that promote its cortical association (Figure 5C and 6C), a construct lacking the C-terminal region of Cyst (Cyst^R^ΔC::GFP) did not rescue the *cyst* RNAi phenotype (Figure 6A,B). However, *cyst* RNAi embryos expressing GFP::Cyst^R^ΔPBM were viable (Figure 6A). This finding indicates that the Cyst C-terminal region, but not the PBM, is essential for epithelial integrity, consistent with our finding that Cyst interacts with PDZ domain-containing proteins Baz and Patj independent of its PBM (see below).

Expression of a construct lacking the Cyst N-terminus (GFP::Cyst^R^ΔN) partially rescued *cyst* shRNA embryos (32% embryonic lethal; Figure 6A). Dead GFP::Cyst^R^ΔN embryos displayed a wild-type looking cuticle, indicating that the N-terminal region of Cyst is dispensable for most of Cyst activity. As Cyst contains a putative aPKC phosphorylation site in its N-terminal region at S320 (Wang et al., 2012a), we asked whether aPKC phosphorylation of Cyst could potentially modify its activity. However, non-phosphorylatable and phosphomimetic isoforms of Cyst (GFP::Cyst^R^S320A and GFP::Cyst^R^S320E) fully rescued *Cyst* shRNA embryos (Figure 6A).

We also tested the impact of Cyst domains in overexpression experiments. We expressed HA-tagged Cyst isoforms (shown in Figure 5C) with the ubiquitous zygotic driver *da-Gal4* or the maternal driver *mat-GAL4*. Only the full-length protein (HA::Cyst) expressed maternally caused substantial embryonic lethality (Figure 1F), whereas overexpression of other constructs had no effect on embryonic viability. In line with data from our rescue analysis, this suggests that Cyst requires both the DH-PH domain and the C-terminal region for protein function. Taken together, our analysis of genomic rescue constructs and overexpression constructs revealed that both the DH-PH domain and the C-terminal region of Cyst harbour crucial activities. These findings support a model where Cyst acts as a RhoGEF whose activity depends on cortical recruitment. While a C-terminal-dependent recruitment mechanism seems particularly important, multiple regions of the Cyst protein can contribute to cortical recruitment or retention at least in part independently.

### Apical recruitment of Cyst requires physical interaction with the Crb complex and Baz/Par3

Previous work indicated that mammalian p114RhoGEF forms complexes with Lulu2 (also known as EPB41L4B, one of two mammalian homologs of Drosophila Yrt), Patj and Par3 (Nakajima and Tanoue, 2011). To probe for similar interaction among Drosophila proteins we first asked whether the Crb complex, which includes Patj, and Baz are required for Cyst recruitment to the apico-lateral cortex. We injected GFP::Cyst expressing embryos with dsRNA against *crb* or *baz*. In both cases, GFP::Cyst was lost from the cortex by stage 9 (Figure 7A).

**Figure 7.**
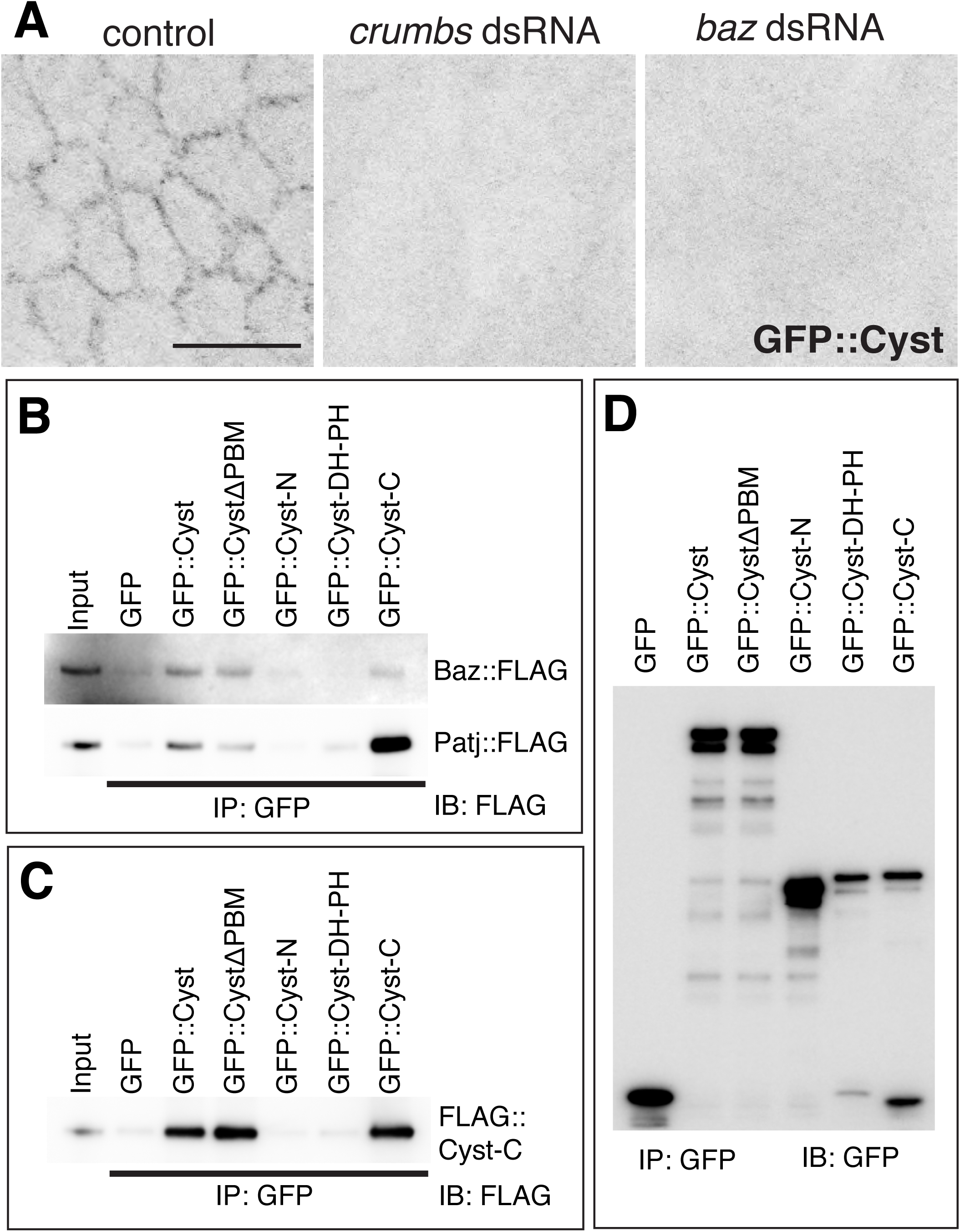
Cyst physically interacts with Baz and the Crb complex. **(A)** Z-projections taken from live stage 9 embryos expressing GFP::Cyst. Embryos have been injected with dsRNAs against *crb* or *baz*. GFP::Cyst is lost from the cortex in knockdown embryos. Scale bar, 10 μm. **(B)** Immunoprecipitation with anti-GFP followed by Immunoblot (IB) with anti-FLAG. Cyst constructs were co-expressed with Baz::FLAG and Patj::FLAG in HEK293T cells. Input shows Baz::FLAG and Patj::FLAG bands. GFP::Cyst and GFP::CystΔPBM formed Baz::FLAG or Patj::FLAG containing complexes. GFP::Cyst-C and Patj::FLAG formed a particularly strong complex. GFP::Cyst-C and Baz::FLAG formed a weak complex. **(C)** Immunoprecipitation with anti-GFP followed by Immunoblot (IB) with anti-FLAG. A FLAG-tagged version of Cyst C-terminal region (FLAG::Cyst-C) was co-expressed with various GFP-tagged fragments of Cyst or GFP as a control in HEK293T cells. Input shows FLAG::Cyst-C. GFP::Cyst, GFP::CystΔPBM, and GFP::Cyst-C formed FLAG::Cyst-C containing complexes. **(D)** Controls for (B) and (C). Immunoprecipitation with anti-GFP followed by Immunoblot with anti-GFP. All GFP-tagged constructs are expressed.

As the Cyst C-terminal region displays strong cortical affinity, we asked whether the PDZ scaffolding proteins Baz or Patj can form a complex with the Cyst C-terminal region. We co-expressed various GFP-tagged fragments of Cyst with FLAG-tagged Baz or Patj (Baz::FLAG or Patj::FLAG) in HEK293T cells. We found that GFP::Cyst and GFP::Cyst-C coimmunoprecipitated with Baz::FLAG or Patj::FLAG, with the interaction between Cyst-C and Patj appearing particularly robust (Figure 7B,D). Interestingly, GFP::CystΔPBM also formed Baz::FLAG or Patj::FLAG containing complexes, suggesting that the interaction between Cyst and Baz or Patj takes place in the C-terminal region of Cyst but does not require the Cyst PBM. To further assess whether Cyst and Baz can interact, we co-expressed FLAG-tagged versions of Cyst N, DH-PH and C-terminal regions with GFP-tagged Par3 (GFP::Par3) in HeLa cells. We found that FLAG::Cyst-C and GFP::Par3 appeared to co-aggregate in puncta (Figure S2). These findings suggest that Drosophila Cyst undergoes molecular interactions with the Crb complex and Baz to support its apico-lateral localization. In contrast to findings in mammalian cells, we did not detect molecular interactions between Yrt and Cyst. This correlates with the observation that Yrt acts as a basolateral polarity protein in early Drosophila embryos and that the *yrt* mutant phenotype is rather different from the *cyst* phenotype (Laprise et al., 2006, 2009).

To further explore the function of the C-terminal region of Cyst which is crucial for Cyst localization we co-expressed FLAG::Cyst-C with various GFP-tagged fragments of Cyst in HEK293T cells. We detected binding of GFP::Cyst, GFP::CystΔPBM, and GFP::Cyst-C to FLAG::Cyst-C (Figure 7C,D). These constructs all contain the Cyst coiled-coil domain (CC), which facilitates oligomerization in some CC proteins (Schultz et al., 1998; Letunic et al., 2015), suggesting that Cyst oligomerization through its C-terminal region may contribute to the cortical affinity associated with this region.

Taken together, our data support the view that Cyst is directed to the apico-lateral cortex through a multifaceted mechanism, involving several membrane-targeting sequences, oligomerization of the Cyst C-terminal region, and redundant and/or parallel interactions between Cyst C-terminal region and Baz and the Crb complex.

### Cyst targets Rho1

RhoGEF domains often show specificity for Rho, Rac, or Cdc42 although there are some examples of promiscuity (McCormack et al., 2013; Ngok et al., 2014). Phylogenetic analysis suggests that Cyst is the single orthologue of a group of four mammalian RhoGEFs that target RhoA in cell culture. To ask whether Cyst targets Rho1 *in vivo*, we assayed the activity of two probes that are thought to preferentially bind to Rho1-GTP, Anillin-RBD::GFP (Munjal et al., 2015) (Figure 8A,B) and PKNG58A::GFP (Simoes et al., 2014) (Figure S3). The junctional localization of both probes was reduced by approximately 30-60% in *cyst* compromised embryos compared to controls.

**Figure 8.**
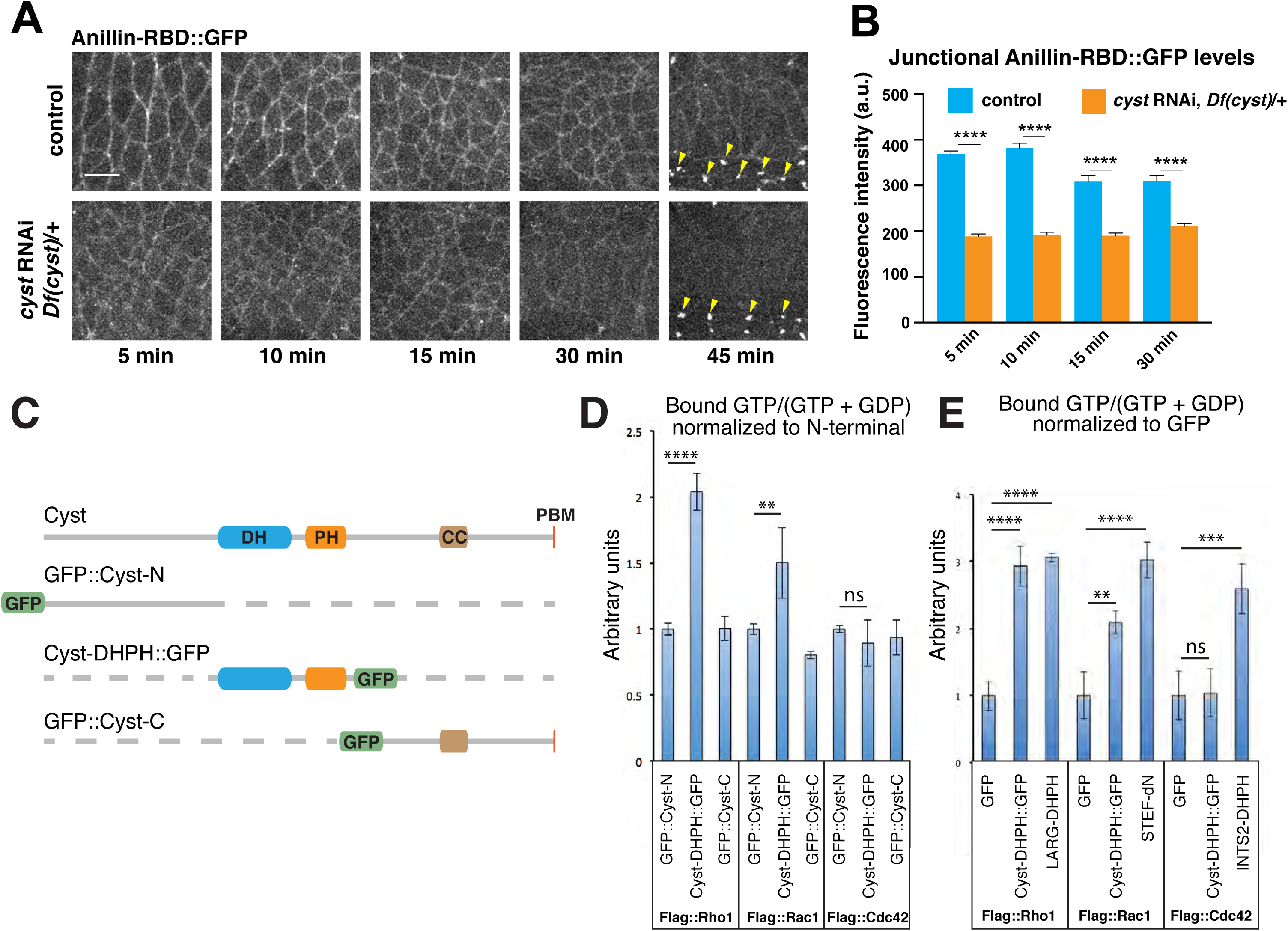
Cyst preferentially targets Rho1. **(A)** Stills of time-lapse movies (Movies S3 and S4) showing the Rho activity probe Anillin-RBD::GFP in a control and in an embryo derived from a *cyst* RNAi, *Df(cysts)*/+ mother. Time is minutes after the onset of germband extension. Junctional levels of Anillin-RBD::GFP are reduced in *cyst* depleted embryos. In ventral midline cells, the probe accumulates also in midbodies after cell division (arrowheads) in control and *cyst* depleted embryos. Scale bar, 10 μm. **(B)** Quantification of junctional levels of Anillin-RBD::GFP in control and *cyst* depleted embryos at the time points indicated in (A). Quantifications at 45 min were not included as the probe’s junctional signal was undetectable in *cyst* depleted embryos. Bars indicate mean +/-SEM. n=4 embryos/condition; 300-387 cell edges analyzed per time point. From left to right: **** p =1.5×10^−60^, 8.3×10^−42^, 1.2×10^−9^, 1.3×10^−10^ (KS test). **(C)** Schematic of GFP-tagged fragments of Cyst used in (D) and (E). **(D)** Activation of Drosophila Rho1, Rac1, or Cdc42 by the indicated fragments of Cyst protein depicted in (C). GTPases were immunoprecipitated from HEK293T cells co-expressing the Cyst fragments. GTP and GDP were eluted and subsequently analyzed by LC-MS/MS. The absolute amount of GTP and GDP are shown as a % normalized to co-expression with GFP::Cyst-N. Cyst-DHPH::GFP shows a 2-fold relative increase in Rho1 associated GTP, and a 1.5-fold relative increase in GTP-associated Rac1. No detectable activation of Cdc42 was found. GFP::Cyst-N and GFP::Cyst-C act as negative controls. Error bars represent SD of four independent experiments. ****, p<0.0001; **, p<0.01; ns, not significant (t-test). **(E)** Positive controls for (D) using mammalian GEF proteins. The DH-PH domain of LARG (LARG-DHPH), an active version of STEF (STEF-ΔN), and the DH-PH domain of INTS2 (INTS2-DHPH) are known to target Rho1, Rac1, and Cdc42 respectively. GFP-DHPH and LARG-DHPH show comparable targeting of Rho1. GFP-DHPH targeting of Rac1 was less than that seen for STEF-ΔN. Values are normalized to GFP, which acts as a negative control. ****, p<0.0001; ***, p<0.001; **, p<0.01; ns, not significant (t-test).

To further assess interactions between Cyst and Rho1, we asked whether Cyst activates Rho1 in cell culture. We co-expressed FLAG-tagged versions of *Drosophila* Rho1, Rac1, and Cdc42 along with the DH-PH domain of Cyst fused to GFP (Cyst-DHPH::GFP) or N- or C- fragments (GFP::Cyst-N or GFP::Cyst-C) as controls in HEK293T cells (Figure 8C). Immunoprecipitation of the GTPases where followed by direct analysis of absolute levels of associated GTP and GDP by LC-MS/MS. Co-expression of Cyst-DHPH::GFP with Rho1 showed a 2-fold increase in Rho1-associated GTP as normalized to the levels seen with GFP::Cyst-N (Figure 8D). As a positive control, the DH-PH domain of LARG, a known mammalian RhoA-GEF (Cook et al., 2014), showed a level of Rho1 GTP loading comparable to Cyst-DHPH::GFP (Figure 8E). These data are consistent with our observation that the expression of Cyst-DHPH::GFP, but not GFP::Cyst-N or GFP::Cyst-C, produces dorsal ruffling and stress fiber formation in HeLa cells (Figure S2), a phenotype reminiscent of RhoA activation in fibroblasts (Hall, 2012; Hanna and El-Sibai, 2013). We detected a 1.5-fold relative increase in GTP-associated Rac1. However, the level of Rac1 targeting by Cyst-DHPH::GFP was less than that seen for an active version of the Rac-specific GEF STEF-ΔN (Matsuo et al., 2003), which acted as a positive control (Figure 8D,E). No detectable activation of Cdc42 was found.

Finally, we tested the ability of Cyst to suppress the effects of dominant-negative (DN) isoforms of Rho1 (Strutt et al., 1997), Rac1, and Cdc42 (Luo et al., 1994). Expression of any DN GTPases under mat-GAL4 control produces a prominent cuticle phenotype. This is presumably due to the sequestration of GEFs preventing activation of Rho, Rac, or Cdc42. We hypothesized therefore that overexpression of a GEF could rescue the effects of a DN GTPase if active levels of its target GTPase are restored. We co-expressed HA::Cyst or the N-terminal region of Cyst, Cyst-N as a negative control, with DN versions of Rho1, Rac1, or Cdc42. Overexpression of Cyst produced a cuticle that is intact with the exception of a head defect (Figure 1F). We found that co-expression of HA::Cyst with DN-Rho1 partially rescues the DN-Rho1 cuticle defects (Figure S3). In contrast, no rescue was observed with Cyst-N. Co-expression of HA::Cyst and DN-Rac1 led to an additive phenotype (Figure S3), and co-expression of HA::Cyst and DN-Cdc42 showed no obvious genetic interaction. Collectively, our genetic and biochemical data support the conclusion that Cyst preferentially activates Rho1.

### Cyst and Crb are required for normal myosin II enrichment at AJs

AJ stability is tightly connected to actomyosin dynamics (Lecuit and Yap, 2015). To ask whether Cyst plays a role in actomyosin dynamics at AJs, we examined the distribution of myosin II by following fluorescent protein-tagged regulatory light chain of non-muscle myosin II (Spaghetti Squash; Royou et al., 1999) in live embryos derived from *cyst* RNAi, *Df(cyst)*/+ mothers (Figure 9A,B,D). In the ectoderm, myosin II is recruited to the apico-lateral cortex during stage 6 and 7 just before the onset of germband extension. Defects in myosin II were noted from stage 7 onwards with junctional myosin levels reduced and less uniformly distributed around the apical cell perimeter. Interestingly, a similar reduction in junctional myosin was observed in *crb* RNAi embryos (Figure 9C,E). In contrast, medial myosin II levels behaved differently in Crb and Cyst depleted embryos as germband extension progressed; whereas medial myosin II levels were enhanced in Crb compromised embryos compared to controls, Cyst depleted embryos showed a moderate reduction in medial myosin II (Figure 9D,E).

**Figure 9.**
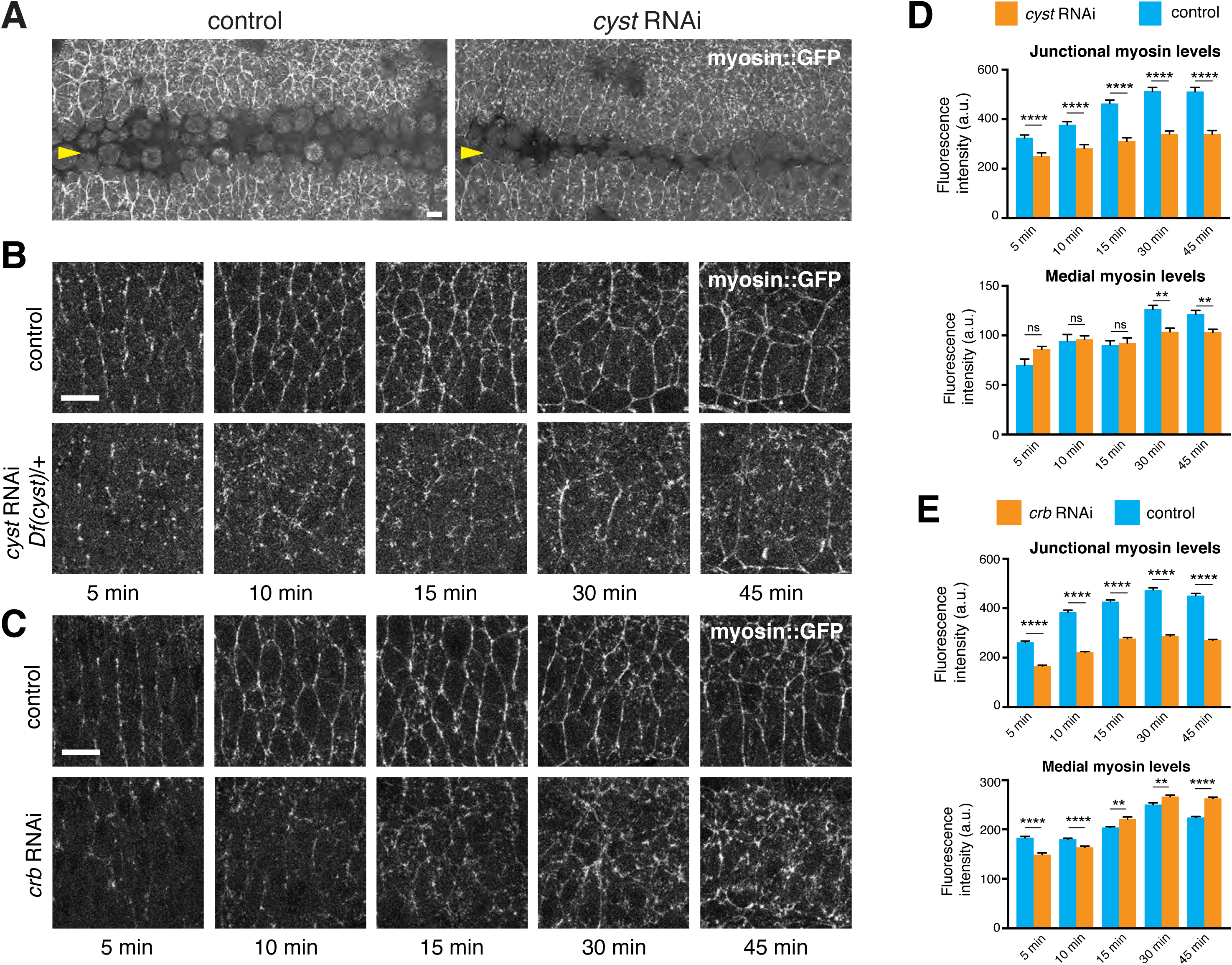
Cyst regulates actomyosin dynamics at AJs. **(A)** Z-projections taken from live stage 8 embryos expressing myosin::GFP (Sqh::GFP) in a wild-type or *cyst* RNAi background. The ventral ectoderm is shown with the arrowhead pointing to ventral midline. *cyst* RNAi embryo shows a reduction of myosin::GFP. Scale bar, 10 μm. **(B)** Stills of time-lapse movies (Movies S5 and S6) showing myosin:GFP in a control embryo (maternal expression of mCherry RNAi) and in a *cyst* depleted embryo (maternal expression of *cyst* RNAi in a *Df(cysts)*/+ mother). Time is minutes after the onset of germband extension. In the control, multicellular myosin II cables form at the anterior-posterior cell-cell contacts during axis elongation. Junctional myosin levels and multicellular cables are reduced in *cyst* depleted embryos. Scale bar, 10 μm. **(C)** Stills of time-lapse movies (Movies S7 and S8) showing myosin:GFP expression in a control embryo (H_2_O injected) and in a *crb* dsRNA injected embryo. Time is minutes after the onset of germband extension. Junctional myosin levels are reduced and medial myosin levels gradually become enhanced in the *crb* depleted embryo. **(D)** Quantification of junctional and medial myosin levels in control and *cyst* depleted embryos at the time points indicated in (B). Bars indicate mean +/- SEM. n=3 embryos/condition; 305-557 cell edges and 357-525 apical surfaces analyzed per time point. For junctional myosin, from left to right: **** p =4.2×10^−4^, 5.9×10^−5^, 3.1×10^−16^, 4.2×10^−21^, 3.4×10^−16^ (KS test). For medial myosin, from left to right: ns, p=0.19; 0.59; 0.99; ** p=6×10^−2^ (KS test). **(E)** Quantification of junctional and medial myosin levels in control and *crb* RNAi embryos at the time points indicated in (C). Bars indicate mean +/- SEM. n=3-5 embryos/condition; 554-1244 cell edges and 580-800 apical surfaces analyzed per time point. For junctional myosin, from left to right: **** p =6.5×10^−53^, 5.5×10^−72^, 2.5×10^−60^, 7.2×10^−66^, 3.0×10^−62^ (KS test). For medial myosin, from left to right: ****, p=1.1×10^−6^; 8.8×10^−8^; **, 2×10^−3^; 1×10^−3^; **** p=2.7×10^−9^ (KS test).

We also found that the loss of Cyst causes aberrant F-actin distribution and dynamics as assayed by Utrophin::GFP (Stage 11/12 - Figure S4), including in small groups of cells that go on to form epithelial cysts Moreover, high levels of F-actin were associated with apical protrusions (Figure S4), consistent with a loss of Rho activity and AJ integrity, which are known to limit protrusive activity mediated by Rac (Harris and Tepass, 2010). Taken together, these findings indicate that Cyst is required for the normal association of actomyosin with apical AJs and support a model posing that Cyst couples the apical Crb complex to junctional Rho1 activity and AJ stability.

## Discussion

### Cyst links epithelial polarity, AJ stability and actomyosin remodelling

Antagonistic interactions between apical and basolateral polarity regulators position AJs at the apico-lateral membrane to form a junctional complex. In turn, AJs are thought to maintain apical-basal polarity through the segregation of the apical and basolateral membrane domains, organization of the cytoskeleton, and direct polarity by acting as signaling centers for polarity complexes (Harris and Tepass 2010; Laprise and Tepass, 2011; Tepass, 2012; Harris, 2012). Although a number of *Drosophila* RhoGEFs and RhoGAPs have been implicated in epithelial polarity and AJ stability (McCormack et al., 2013; Mack and Georgiou, 2014), no single RhoGEF or RhoGAP has been found to phenocopy the polarity or junctional defects that are seen in embryos compromised for factors like Crb, aPKC, or E-cadherin (Tepass and Knust, 1990; Tepass et al. 1996; Hutterer et al., 2004). Our findings suggest that loss of the RhoGEF Cyst causes polarity phenotype strikingly similar to the loss of core apical polarity proteins such as Crb. Moreover, we find that Cyst is recruited to the apico-lateral cortex by the action of polarity proteins and by activating Rho1 stabilizes AJs-associated actomyosin to support junctional and epithelial integrity.

In *Cyst* compromised embryos, AJ formation is disrupted in early gastrulation and AJs do not form a circumferential belt. These defects in AJ assembly or stability correlate with reduced and irregular myosin accumulation at the apico-lateral cortex. Given the molecular function of Cyst as a GEF for Rho1, loss of myosin activity is presumably the immediate cause for the defects in AJ formation, and the subsequent loss of apicobasal polarity in many epithelial cells. Also *crb* mutants showed a similar decline in junctional myosin. Therefore, a major function of the apical Crb polarity complex appears to be the Cyst-mediated support of junctional actomyosin (Figure 10).

**Figure 10:**
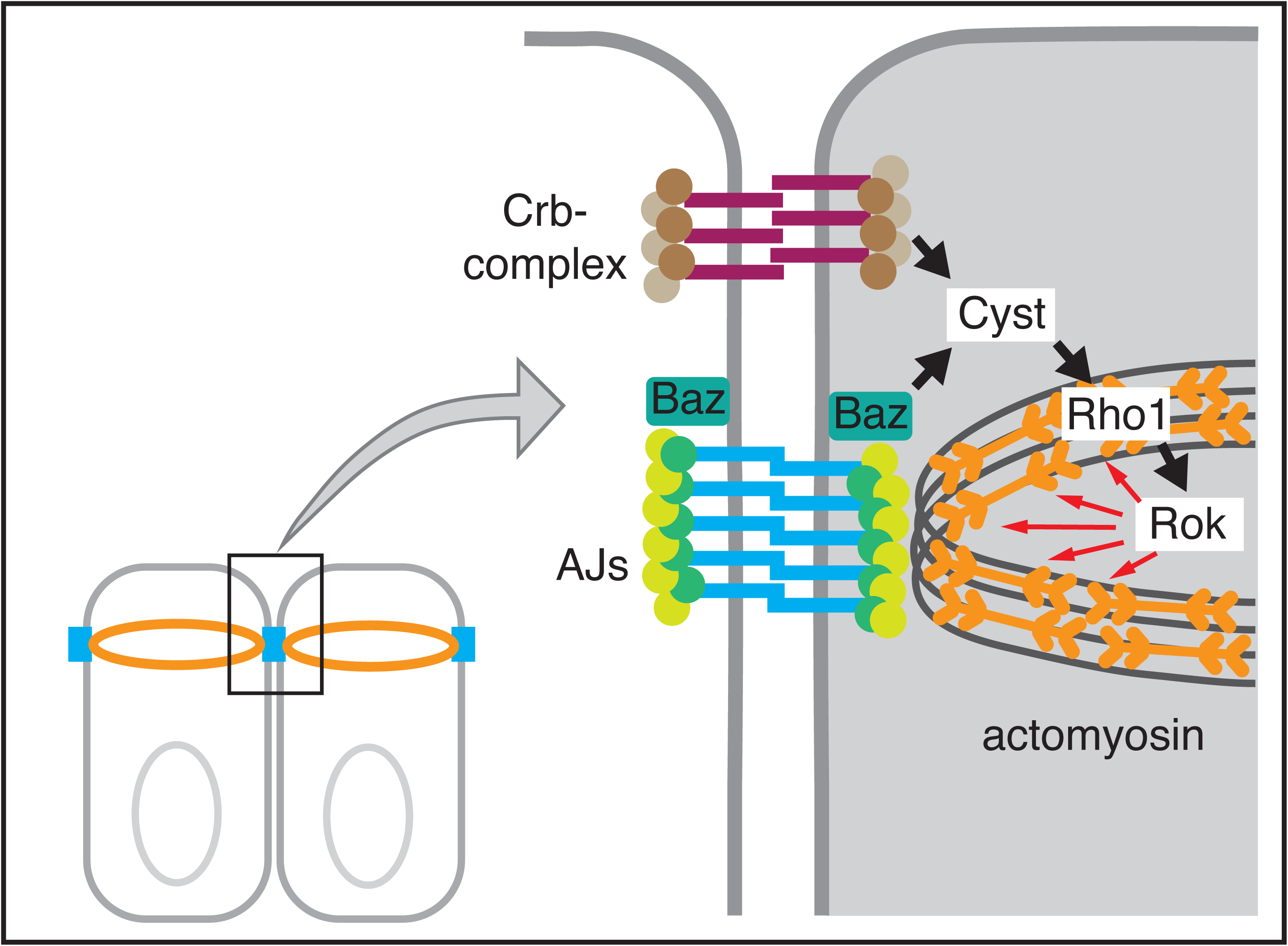
Model of Cyst function in the Drosophila embryo. The Crb complex and Baz recruit the RhoGEF Cyst to the apicolateral membrane where it activates Rho1 and myosin to support of junctional and epithelial integrity.

While many cells in *crb* or *cyst* mutants undergo programmed cell death, others retain or recover polarity and form small epithelial cysts, a process seen from mid-embryogenesis (post-gastrulation stages) onward. Several polarity proteins such as Crb, Sdt and Baz are needed for normal epithelial polarization in early embryos but are not essential for polarization in post-gastrulation embryos, which explains the ability of some epithelial cells in these mutants to form epithelial cysts with normal polarization (Tanentzapf and Tepass, 2003; Bilder et al., 2003; Laprise et al., 2009). In fact, when programmed cell death is suppressed, cyst formation is shown by all epithelial cells in *crb* mutants (Tanentzapf and Tepass, 2003). Formation of epithelial cysts seen in *cyst* mutant embryos therefore suggests that Cyst is also not essential for epithelial polarity in late embryos. This view is supported by the steady decline of Cyst protein accumulation at AJs seen in late embryos.

Several observations including genetic interaction of *cyst* with genes encoding basolateral polarity proteins, the dependency of the junctional localization of Cyst on the apical polarity proteins Baz and Crb, the physical interactions between Cyst and apical polarity proteins, and the function of Cyst in stabilizing AJs, indicate that Cyst is an integral part of the apical polarity machinery in early Drosophila embryos. A particular striking finding was the complete suppression of the cyst phenotype by co-depletion of the basolateral polarity proteins Scrib or Lgl in that double mutant embryos showed phenotypes indistinguishable from single *scrib* or *lgl* mutant embryos. This mimics previous observations with double mutants of *crb* or *sdt* and *scrib*, *lgl* or *discs large* (Tanentzapf and Tepass, 2003; Bilder et al., 2003). Moreover, we found that a reduction of aPKC enhanced Baz mislocalization in Cyst compromised embryos suggesting that aPKC cooperates with Cyst and acts upstream or in parallel to Cyst to organize Baz. These findings emphasize that Cyst like Crb and aPKC is a component of a negative feedback circuit between apical and basolateral regulatory networks that govern epithelial polarity. The dependency of Cyst localization on Crb and Baz suggests that Cyst acts downstream of these two proteins. Once polarized, Cyst appears to maintain polarity and junctional stability through actomyosin remodelling.

### Cyst associates with the cortex through a multifaceted mechanism

Our results suggest that Cyst has multiple membrane targeting sequences. The N, DH-PH, or C– terminal regions of Cyst expressed on their own were all found to associate with the cortex at the level of AJs. Of these, the Cyst C-terminal region demonstrated particularly strong cortical affinity, as CystΔC was only detected at the cortex in *cyst* depleted embryos but not in the presence of endogenous Cyst. Moreover, our *in vivo* structure-function data indicate that the C-terminal region is essential for Cyst activity. We propose therefore that C-terminal-dependent cortical association of Cyst is crucial for protein function. Interestingly, we found that Cyst can oligomerize through its C-terminus which includes a CC domain that could facilitate oligomerization (Schultz et al., 1998; Letunic et al., 2015). Clustering of Cyst could enhance its cortical association.

Patj represents one possible anchor for Cyst clusters at the cortex. Our biochemical data show that the Cyst C-terminal region is sufficient for interactions with the scaffolding protein Patj, which couples Cyst to the Crb-complex. Patj has been implicated as a myosin II activator in the embryo (Sen et al., 2012). Given our genetic and biochemical data it is likely that Crb, Patj and Cyst form a complex that organizes junctional actomyosin. However, as Patj is not essential for embryonic survival (Sen et al., 2012), Cyst may interact with additional binding partners within the Crb complex. Another apical binding partner for Cyst is Baz/Par3, a conclusion supported by several lines of evidence. The Cyst C-terminal region co-precipitates with Baz when co-expressed in HEK293T cells and Cyst co-aggregates with GFP::Par3 in HeLa cells. Moreover, we found that Cyst was lost from the cortex in embryos depleted of Baz. Taken together, our data support a model in which Cyst is directed to the apico-lateral cortex through a mechanism involving multiple membrane targeting sequences, the oligomerization of Cyst C-terminal region, and interactions with Baz and the Crb complex.

### The GEF activity of Cyst targets Rho1

We found that Cyst becomes enriched at the apico-lateral cortex after the mesoderm and endoderm have invaginated and the germband starts to elongate. This localization coincides with the assembly of the apical-cortical actomyosin network. Rho-Rok signalling plays a critical role in the activation of myosin II in this process (Lecuit and Lenne, 2007; Amano et al., 2010; Harris and Tepass, 2010). Our structure-function analysis showed that Cyst contains an essential RhoGEF domain as predicted, and the use of Rho activity probes, genetic interaction, and biochemical data showed that Cyst preferentially targets Rho1. We propose therefore that Cyst activates Rho1 to organize actomyosin at the cortex at a time when AJs assemble into a circumferential belt (stage 6/7). Consistent with this, we found that Cyst is important for maintaining normal cortical levels of myosin II. A similar loss in junctional myosin was also observed in Crb compromised embryos in line with our finding that Crb is required for Cyst junctional recruitment. The *cyst* mutant phenotype suggests that Cyst is the key RhoGEF that activates Rho1 at ectodermal AJs. In contrast, RhoGEF2 functions in the mesoderm and ectoderm where it becomes apico-cortically enriched and activates Rho1 to recruit myosin II to the apical-medial cortex (Barmchi et al., 2005; Manning and Rogers, 2014; Kerridge et al., 2016). Thus, RhoGEF2, Cyst, and potentially other RhoGEFs act together on Rho1 to orchestrate the balance of cortical and medial myosin dynamics.

### Cyst and its mammalian orthologs share a conserved function

Cyst is the single orthologue of a group of four mammalian RhoGEFs that target RhoA in cell culture (Cook et al., 2014). One of the mammalian orthologs, p114RhoGEF stabilizes tight junctions and AJs through the organization of actin cytoskeleton associated with cellular junctions (Nakajima and Tanoue, 2010; 2011; Terry et al., 2011; Acharya et al., 2018). p114RhoGEF is recruited to apical junctions through a mechanism involving Par3 and Patj (Nakajima and Tanoue, 2011) and the heterotrimeric G protein Gα12 and the G protein-coupled receptor (GPCR) Sphingosine-1 phosphate receptor 2 (Acharya et al., 2018). G proteins and GPCRs have also been linked to actomyosin in the Drosophila ectoderm in a pathway mediated by RhoGEF2 (Kerridge et al., 2016). Thus, it is possible that also Cyst is controlled by a GRCR pathway in addition to its dependency on the Crb complex and Baz. The C-terminal PBM of p114RhoGEF appears to be the minimal region required for cortical association (Nakajima and Tanoue, 2011). Although the putative Cyst C-terminal PBM is required for normal cortical localization of Cyst, we found that it was dispensable for protein function or complex formation with Baz or Patj. Further, p114RhoGEF requires the polarity regulator Lulu2 (a homolog of Drosophila Yrt) to activate RhoA (Nakajima and Tanoue, 2010; 2011). In contrast, we did not detect genetic or biochemical interactions between Cyst and Yrt in Drosophila. Recently, *ARHGEF18*, the human ortholog of *p114RhoGEF*, was identified as a gene associated with retinal degeneration (Arno et al., 2017), and a fish ortholog is required to maintain epithelial integrity of the retina (Herder et al., 2013). *ARHGEF18* mutant retinal defects closely resemble those found in patients carrying mutations in the *crb* homolog *CRB1* (Arno et al., 2017). We conclude that the function of Cyst and p114RhoGEF/ARHGEF18 in coupling apical polarity proteins to junctional Rho activity and actomyosin function is conserved between flies and mammals and likely contributes to retinal health in humans, although some of the molecular interactions may have shifted in relative importance.

The other mammalian orthologues of Cyst, p190RhoGEF, AKAP-13, and GEF-H1 have not been implicated as regulators of epithelial polarity (Cook et al. 2014). GEF-H1 (aka ARHGEF2 and Lfc) was shown to be inactive at mature tight junctions (Aijaz et al., 2005; Terry et al., 2011). In this case, the tight junction protein Cingulin forms a complex with GEF-H1, preventing it from activating RhoA (Aijaz et al., 2005; Terry et al., 2011). Instead, GEF-H1 is thought to promote junction disassembly and cell proliferation, presumably through an association with the mitotic spindle (Ren et al., 1998; Aijaz et al., 2005; Samarin et al., 2007; Birkenfeld et al., 2008; Terry et al., 2011; Cullis et al. 2014). GEF-H1 was also implicated in the morphogenesis of the vertebrate neural tube (Itoh et al., 2014), and in the regulation of RhoA activity during cytokinesis (Birkenfeld et al., 2007). Like GEF-H1, p190RhoGEF has been shown to associate with microtubules (Birkenfeld et al., 2008). GEF-H1 and AKAP-13 were also found to serve additional functions independent of their RhoGEF activity (Shibolet et al., 2007; Cullis et al., 2014). Whether and how Cyst might consolidate the functions of its various mammalian orthologues remains to be explored.

## Materials and Methods

### Generation of the *cyst^1^* mutation

We generated a null mutant for *cyst* (*cyst^1^*) using the RNA-guided CRISPR/Cas9 system (Gratz et al., 2013; GenetiVision Corporation, TX, US). The second exon of *cyst* was targeted using gRNA1 (GTTAGCAATAACTAATCGCA) and gRNA2 (AGCTCCTCGAGCCAAGCCCG) and replaced with a 3xP3-GFP cassette. Sequencing confirmed the following breakpoints: 19503155 and 19506410.

### Drosophila genetics

Flies were raised on standard media at 25°C. Cyst was depleted in females carrying mat-GAL4 (*P{matα4-GAL-VP16}67; P{matα4-GAL-VP16}15*; Häcker and Perrimon, 1998) and *cyst* RNAi (Valium20-SH00146.N-40 inserted at attP40 on the 2^nd^ chromosome; Ni et al., 2011; Transgenic RNAi Project [TRiP]). Virgin females were crossed to males carrying *cyst* RNAi to produce cyst depleted embryos. A second insertion of the same *cyst* shRNA (Valium22-SH00146.N2) gave similar results. UAS-constructs expression in Figure 4 was facilitated by a maternal-triple-driver (MTD-GAL4; BDSC #31777).

The following fly lines were used: *cyst^1^* FRT40A/ CyO (this work), *cyst* RNAi (Valium20-SH00146.N-40, BDSC #38292), *Df(cyst)* (*Df(2L)BSC301*, BDSC #23684), *Df(cyst)* (*Df(2L)BSC301*); *cyst* RNAi (Valium22-SH00146.N2, BDSC #41578), *scrib* RNAi on 3R (Valium20-SH02077.N BDSC #35748), *yw*; *lgl^4^ FRT40A* (gift from T. Xu, Yale University, USA). yw; *lgl^4^*, *cyst*^1^ FRT40A, *aPKC^K06403^* (gift of C. Doe, University of Oregon, USA), *mCherry* shRNA (BDSC #35785), UAS-PKNG58A::Venus (Simoes et al., 2014), UAS-Anillin-RBD::GFP (Munjal et al., 2015), and *histone::*GFP (gift of A. Wilde, University of Toronto, Canada).

*shg>DEcad::GFP* (Huang et al., 2009) and *sqh>sqh::GFP* (Royou et al., 1999) were recombined with the mat-GAL4 driver. *Utrophin::GFP*; *cyst* RNAi males were crossed to the mat-GAL4 driver, and then crossed to *cyst* RNAi males to generate heterozygotes for *Utrophin::GFP* (eGFP fused to the actin binding domain of human Utrophin; Rauzi et al., 2010). Cyst overexpression constructs were driven by *da*-GAL4 (Wodarz et al., 1995) or mat-GAL4. Females expressing HA::Cyst or HA::Cyst-N by the mat-GAL4 driver were crossed to *P{UAS-Rho1.N19}1.3* (Strutt et al., 1997), *P{UAS-Rac1.L89.}6*, or *P{UAS-Cdc42.N17}* (Luo et al., 1994) males.

### Cuticle Preps of Drosophila embryos

The cuticles of embryos were prepared according to Wieschaus and Nüsslein-Volhard (1986).

### Immunofluorescence

*Drosophila* embryos were heat-fixed unless otherwise stated (Tepass, 1996). To visualize the PKNG58A::Venus sensor, embryos were fixed in a 1:1 mixture of 3.7% formaldehyde in phosphate buffer, pH 7.4, and heptane for 40 min under agitation followed by hand devitalization. Primary antibodies used were: anti-Crb (rat polyclonal, extracellular F3; 1:1000; Pellikka et al., 2002), anti-Yrt (guinea pig polyclonal, GP7; 1:500; Laprise et al., 2006), anti-Arm (mouse monoclonal, N2-7A1; 1:50; Developmental Studies Hybridoma Band[DSHB]), anti-HA (rat monoclonal, 3F10; 1:500; Roche). Anti-Baz (1:3500), anti-Baz (GP, 1:500, a gift from Jennifer Zallen), anti-PKCζ (C-20; 1:100; rabbit polyclonal; Santa Cruz), and anti-Dlg (1:100; mouse; DSHB). For immunostaining embryos were fixed for 20 min in 3.7% formaldehyde diluted in 1:1 PBS:Heptane. Transfected HeLa cells were fixed with 3.7% formaldehyde for 10 min at room temperature, followed by treatment with 0.2% TritonX-100 in PBS with 2 mg/ml BSA for 10 min. Markers were mouse anti-FLAG (Sigma), anti-GFP (Abcam; chick), and phalloidin (Invitrogen). All secondaries were IgG (H+L) Alexa Fluor® antibodies (Thermo Fisher Scientific).

### Imaging and quantifying signal intensity

All scanning confocal microscopy was conducted using a Leica TCS SP8 with 40X, 63X, and 100X objectives (HC PL APO CS2 1.30, 1.40, and 1.40 respectively). Time-lapse acquisitions were done in a similar manner as previously described (Blankenship et al., 2006). 3-5 live embryos were examined for each genotype. A 0.5 μm step was used to collect z-stacks. Stills and movies were assembled from maximum-intensity projections of 6 apical planes (ImageJ). Adobe Photoshop and Adobe Illustrator were used to process and arrange images. The same settings were applied to all images within an experimental series. The average fluorescence intensity of junctional and medial Sqh::GFP, Anillin-RBD::GFP, and PKNG58A::Venus was quantified in segmented cells using MATLAB and the script SIESTA (scientific image segmentation and analysis; Fernandez-Gonzalez and Zallen, 2011). To quantify junctional Sqh::GFP, Anillin-RBD::GFP, and PKNG58A::Venus, we manually drew 3-pixel-wide lines (180 nm/pixel) for cell edges at the indicated time points to obtain the mean pixel intensity, and the mode (cytoplasmic) intensity was subtracted for background correction. To quantify medial Sqh::GFP mean protein levels, each cell was divided into two compartments (Fernandez-Gonzalez and Zallen, 2011). The junctional compartment was determined by a 3-pixel-wide (0.54 µm) dilation of the cell outline identified using watershed or LiveWire segmentation in SIESTA. The medial compartment was obtained by inverting a binary image representing the junctional compartment. Given the dynamic nature of medial Sqh::GFP, we quantified it as the mean pixel intensity in the medial compartment considering 10 consecutive time frames per cell (total of 5 min), centered at the indicated time points, and subtracted the mode (cytoplasmic) intensity for background correction.

In Figure 4, spinning disk confocal microscope (Quorum Technologies) with 63X Plan Apochromat NA 1.4 (Carl Zeiss) objective, piezo top plate, and EM CCD camera (Hamamatsu Photonics) was used. Baz puncta were quantified by dividing the number of cells with two Baz puncta by the total number of cells. aPKC levels were the average difference between the cell cortical and cytoplasmic signals of five different cells per embryo, and normalized to internal GFP-expressing controls. Cytochalasin D and DMSO were diluted 2000 times in an NaCl-octane solution prior to embryo incubations.

## Molecular Biology

### Genomic rescue constructs

A ~7.0 kb fragment (genomic region 19509164 – 19502181) encompassing the *cyst* gene was amplified from BACR27M12 (BACPAC) and recombined into pENTR221 (Invitrogen). To confer RNAi-resistance on the resulting Entry Clone, silent mutations were introduced at two distinct sites corresponding to SH00146.N and SH04062.N (TRiP; the latter shRNA subsequently proved ineffective). All other genomic constructs were derived from this Entry Clone. The GFP-tagged full-length construct was generated by inserting sfGFP (Drosophila codon-optimized superfolder GFP; Pédelacq et al., 2006) in between the ATG and the second codon of *cyst*. The GFP-tagged deletion constructs were generated by replacing the domains described in Figure 6A with sfGFP. In all constructs an S(GGGGS)_2_ linker was introduced in between GFP and Cyst, and in CystΔN the intron was left intact to preserve potential regulatory sequences. Finally, all Entry Clones were recombined into the Drosophila transformation vector pBID-G (Wang et al., 2012b).

### UAS constructs

The *cyst* ORF including the intron was amplified from the first Entry Clone described above and recombined into pDONR221. The intron was then deleted from this Entry Clone to give the full-length *cyst* ORF. Fragments corresponding to the domains described in Figure 5C were amplified from this Entry Clone and recombined into pDONR221. The resulting Entry Clones were recombined into a destination vector containing the *UASp* promoter (Rørth, 1998) and either an N- or C-terminal 3×HA tag separated from Cyst by an SGGGS linker (F. Wirtz-Peitz; unpublished). GFP and FLAG–tagged Cyst constructs were generated in a similar manner.

All cloning was performed using Gateway (Invitrogen) or In-Fusion (Clontech) cloning kits. All plasmids generated by PCR were sequence-verified along the entire length of the insert. Plasmid and primer sequences are available upon request. Transgenes were placed in attP40 by ΦC31-mediated transgenesis. Injection of dsRNA has been previously described for *baz* (Simões et al., 2010). The dsRNA against *crb* was *in vitro* transcribed from a template that was PCR amplified from genomic DNA using the following *crb* primers fused to a T7 promoter sequence at the 5’ end: forward (5’>3’) CGAGCCATGTCGGAATGGATCAACC; reverse GTCGCTCTTCCGGCGGTGGCTTCAG.

### Cell culture and immunoprecipitation LC-MS/MS

HeLa cells or HEK293T cells were cultured in DMEM with 10% FBS, penicillin, and streptomycin, and transfection was performed using polyethyleneimine (Polysciences). Cells were transfected in 10 cm dishes and incubated with serum-free DMEM for 4 h before harvest. For the LC-MS/MS assay, HEK293T cells were co-transfected with FLAG-tagged versions of Drosophila Rho1, Rac1, or Cdc42 and either: shown GFP-tagged Cyst fragments (Figure 7B) or GFP-tagged versions of the mammalian GEFs LARG-DH-PH, STEF-ΔN, or INTS2-DH-PH, or GFP alone. For immunoprecipitation-Western blot experiments, cells were co-transfected with GFP-tagged Cyst fragments and FLAG::C (as described above) or FLAG-tagged Drosophila Baz or Patj. In HeLa experiments, cells were co-transfected with FLAG-tagged Cyst fragments (as described above) and GFP::Par3. Cells were rinsed once with ice-cold PBS and extracted with ice-cold lysis buffer B (10 mM Tris/HCl at pH7.5, 300 mM NaCl, 10 mM MgCl_2_, 1% Triton X-100, 50 μg/ml PMSF, Complete Protease Inhibitor Cocktail (Roche)) and cleared by centrifugation at 15,000 rpm for 10 min at 4°C. For immunoprecipitation, primary antibodies were added to lysates and incubated with rotation for 2 h at 4°C.

For the LC-MS/MS assay, cell lysates were incubated with FLAG M2 Agarose (Sigma) to precipitate the FLAG-tagged GTPases for 2 h at 4°C. The beads were washed three times with lysis buffer B and further washed two times with buffer C (10 mM ammonium bicarbonate, 5 mM MgCl_2_). The bound nucleotides were eluted with 80 μl methanol and 80 μl chloroform followed by shaking for 30 min at room temperature. 80 μl of water was added, and the supernatants were collected after centrifugation at 10,000 rpm for 10 min, followed by vacuum centrifuge drying. The pellets were re-suspended in 15 μl of 20 mM ammonium bicarbonate and guanine nucleotides were quantified by LC-MS/MS.

The chromatographic separation was performed on an ACQUITY HSS T3 column (100 mm x 2.1 mm, 1.8 μm particles, Waters) at 30°C under isocratic conditions (0.3 mL/min, 10 mM ammonium bicarbonate) using an Acquity UPLC H-Class System (Waters). Mass spectrometric analysis was performed using a Xevo TQD triple quadrupole mass spectrometer (Waters) coupled with an electro-spray ionization source in the positive ion mode. The MRM transitions of *m/z* 524 -> 152.1 and *m/z* 444.1 -> 152.1 were used to quantify GTP and GDP, respectively. Sample concentrations were calculated from the standard curve obtained from serial dilution of each nucleotide standard (Sigma-Aldrich). Analytical conditions were optimized using standards solution. The percent of GTP-bound small GTPases was expressed as GTP/(GTP + GDP) x 100%, which normalizes for the amount of immunoprecipitated GTPase protein in each sample. Values were further normalized to that of the Cyst N-terminal fragment or GFP controls. Error bars represent the standard deviation of four independent experiments.

## Acknowledgments

We thank the TRiP at Harvard Medical School (NIH/NIGMS R01-GM084947) for providing transgenic RNAi fly stocks and/or plasmid vectors used in this study. M. Hoshino, K. Kaibuchi and S. Ohno for providing plasmid vectors. Bloomington Drosophila Stock Center for reagents. Henry Hong for technical assistance. J.T.S. was supported by NSERC PGS D and by the Yoshio Masui Prize in Developmental, Molecular and Cellular Biology. F.W.P. was supported by a postdoctoral fellowship from the Human Frontiers Science Program. D.Y. was supported by a Damon Runyon Cancer Research fellowship. Y.L. was supported by NSERC PGS M. This work was supported by NIH/NIGMS R01-GM084947 and NIH/NIGMS R01-GM067761 to N.P. N.P. is an investigator at the Howard Hughes Medical Institute. The work was also supported by a grant from the Canadian Institutes for Health Research to U.T., and by a grant from the Canadian Institutes of Health Research to T.J.C.H. U.T. is a Canada Research Chair in Epithelial Polarity and Development.

## References

Acharya, B.R., A. Nestor-Bergmann, X. Liang, S. Gupta, K. Duszyc, E. Gauquelin, G.A. Gomez, S. Budnar, P. Marcq, O.E. Jensen, Z. Bryant, and A.S. Yap. 2018. A mechanosensitive RhoA pathway that protects epithelia against acute tensile stress. Dev Cell. 47:439–452.e436.

Aijaz, S., D’Atri, F., Citi, S., Balda, M.S. and Matter, K. 2005. Binding of GEF-H1 to the tight junction-associated adaptor cingulin results in inhibition of Rho signaling and G1/S phase transition. Dev. Cell. 8:777–786.

Amano, M., Nakayama, M. and Kaibuchi, K. 2010. Rho-kinase/ROCK: a key regulator of the cytoskeleton and cell polarity. Cytoskeleton. 67:545–554.

Arno, G., Carss, K.J., Hull, S., Zihni, C., Robson, A.G., Fiorentino, A., Black, G., Hall, G., Ingram, S., Gillespie, R. and Manson, F. 2017. Biallelic mutation of ARHGEF18, involved in the determination of epithelial apicobasal polarity, causes adult-onset retinal degeneration. Am J Hum Genet. 100:334–342.

Aspenström, P. 1999. Effectors for the rho GTPases. Curr Opin Cell Biol. 11:95–102.

Barmchi, M.P., Rogers, S. and Häcker, U. 2005. DRhoGEF2 regulates actin organization and contractility in the Drosophila blastoderm embryo. J Cell Biol. 168:575–585.

Benton, R., and D. St Johnston. 2003. Drosophila PAR-1 and 14-3-3 inhibit Bazooka/PAR-3 to establish complementary cortical domains in polarized cells. Cell. 115:691–704.

Bilder, D., M. Schober, and N. Perrimon. 2003. Integrated activity of PDZ protein complexes regulates epithelial polarity. Nat Cell Biol. 5:53–58.

Birkenfeld, J., Nalbant, P., Bohl, B.P., Pertz, O., Hahn, K.M. and Bokoch, G.M. 2007. GEF-H1 modulates localized RhoA activation during cytokinesis under the control of mitotic kinases. Dev Cell. 12:699–712.

Birkenfeld, J., Nalbant, P., Yoon, S.H. and Bokoch, G.M. 2008. Cellular functions of GEF-H1, a microtubule-regulated Rho-GEF: is altered GEF-H1 activity a crucial determinant of disease pathogenesis? Trends Cell Biol. 18:210–219.

Blankenship, J.T., Backovic, S.T., Sanny, J.S., Weitz, O. and Zallen, J.A. 2006. Multicellular rosette formation links planar cell polarity to tissue morphogenesis. Dev Cell. 11:459–470.

Chartier, F.J.M., Hardy, É.J.L. and Laprise, P. 2011. Crumbs controls epithelial integrity by inhibiting Rac1 and PI3K. J Cell Sci. 124:3393–3398.

Cook, D.R., Rossman, K.L. and Der, C.J. 2014. Rho guanine nucleotide exchange factors: regulators of Rho GTPase activity in development and disease. Oncogene. 33:4021.

Cox, R.T., C. Kirkpatrick, and M. Peifer. 1996. Armadillo is required for adherens junction assembly, cell polarity, and morphogenesis during Drosophila embryogenesis. J Cell Biol. 134:133–148.

Cullis, J., Meiri, D., Sandi, M.J., Radulovich, N., Kent, O.A., Medrano, M., Mokady, D., Normand, J., Larose, J., Marcotte, R. and Marshall, C.B. 2014. The RhoGEF GEF-H1 is required for oncogenic RAS signaling via KSR-1. Cancer Cell. 25:181–195.

Fernandez-Gonzalez, R. and Zallen, J.A. 2011. Oscillatory behaviors and hierarchical assembly of contractile structures in intercalating cells. Physical Biology. 8:045005.

Fletcher, G.C., E.P. Lucas, R. Brain, A. Tournier, and B.J. Thompson. 2012. Positive feedback and mutual antagonism combine to polarize Crumbs in the Drosophila follicle cell epithelium. Curr Biol. 22:1116–1122.

Fox, D.T., Homem, C.C., Myster, S.H., Wang, F., Bain, E.E. and Peifer, M. 2005. Rho1 regulates Drosophila adherens junctions independently of p120ctn. Development. 132:4819–4831.

Gamblin, C.L., E.J. Hardy, F.J. Chartier, N. Bisson, and P. Laprise. 2014. A bidirectional antagonism between aPKC and Yurt regulates epithelial cell polarity. J Cell Biol. 204:487–495.

Genova, J.L., Jong, S., Camp, J.T. and Fehon, R.G. 2000. Functional analysis of Cdc42 in actin filament assembly, epithelial morphogenesis, and cell signaling during Drosophila development. Dev Biol. 221:181–194.

Greenberg, L. and Hatini, V. 2011. Systematic expression and loss-of-function analysis defines spatially restricted requirements for Drosophila RhoGEFs and RhoGAPs in leg morphogenesis. Mechanisms of Development. 128:5–17.

Häcker, U. and Perrimon, N. 1998. DRhoGEF2 encodes a member of the Dbl family of oncogenes and controls cell shape changes during gastrulation in Drosophila. Genes & development. 12:274–284.

Hakeda-Suzuki, S., Ng, J., Tzu, J., Dietzl, G., Sun, Y., Harms, M., Nardine, T., Luo, L. and Dickson, B.J. 2002. Rac function and regulation during Drosophila development. Nature. 416:438.

Hall, A. 2012. Rho family GTPases. Biochem Soc Trans. 40:1378.

Hanna, S. and El-Sibai, M. 2013. Signaling networks of Rho GTPases in cell motility. Cellular Signalling. 25:1955–1961.

Harden, N. 2002. Signaling pathways directing the movement and fusion of epithelial sheets: lessons from dorsal closure in Drosophila. Differentiation. 70:181–203.

Harris, T.J., and M. Peifer. 2007. aPKC controls microtubule organization to balance adherens junction symmetry and planar polarity during development. Dev Cell. 12:727–738.

Harris, K.P. and Tepass, U. 2008. Cdc42 and Par proteins stabilize dynamic adherens junctions in the Drosophila neuroectoderm through regulation of apical endocytosis. J Cell Biol. 183:1129–1143.

Harris, T. 2012. Adherens Junctions: From Molecular Mechanisms to Tissue Development and Disease. Springer Science & Business Media. 60.

Harris, T.J. and Peifer, M. 2007. aPKC controls microtubule organization to balance adherens junction symmetry and planar polarity during development. Dev Cell. 12: 727–738.

Harris, T.J. and Tepass, U. 2010. Adherens junctions: from molecules to morphogenesis. Nat Rev Mol Cell Biol. 11:502.

Herder, C., Swiercz, J.M., Müller, C., Peravali, R., Quiring, R., Offermanns, S., Wittbrodt, J., Loosli, F. 2013. ArhGEF18 regulates RhoA-Rock2 signaling to maintain neuro-epithelial apico-basal polarity and proliferation. Development 140:2787–97.

Huang, J., Zhou, W., Dong, W., Watson, A.M. and Hong, Y. 2009. Directed, efficient, and versatile modifications of the Drosophila genome by genomic engineering. Proc Natl Acad Sci. 106:8284–8289.

Hutterer, A., Betschinger, J., Petronczki, M. and Knoblich, J.A. 2004. Sequential roles of Cdc42, Par-6, aPKC, and Lgl in the establishment of epithelial polarity during Drosophila embryogenesis. Dev Cell. 6:845–854.

Itoh, K., Ossipova, O. and Sokol, S.Y. 2014. GEF-H1 functions in apical constriction and cell intercalations and is essential for vertebrate neural tube closure. J Cell Sci. 127:2542–2553.

Jaffe, A.B. and Hall, A. 2005. Rho GTPases: biochemistry and biology. Annu. Rev Cell Dev Biol. 21:247–269.

Jiang, T., McKinley, R.A., McGill, M.A., Angers, S. and Harris, T.J. 2015. A Par-1-Par-3-centrosome cell polarity pathway and its tuning for isotropic cell adhesion. Curr Biol. 25:2701–2708.

Kerridge, S., Munjal, A., Philippe, J.M., Jha, A., De Las Bayonas, A.G., Saurin, A.J. and Lecuit, 2016. Modular activation of Rho1 by GPCR signalling imparts polarized myosin II activation during morphogenesis. Nat Cell Biol. 18:261.

Kolahgar, G., P.L. Bardet, P.F. Langton, C. Alexandre, and J.P. Vincent. 2011. Apical deficiency triggers JNK-dependent apoptosis in the embryonic epidermis of Drosophila. Development. 138:3021–3031.

Laprise, P., Lau, K.M., Harris, K.P., Silva-Gagliardi, N.F., Paul, S.M., Beronja, S., Beitel, G.J., McGlade, C.J. and Tepass, U. 2009. Yurt, Coracle, Neurexin IV and the Na+, K+-ATPase form a novel group of epithelial polarity proteins. Nature. 459:1141.

Laprise, P. and Tepass, U. 2011. Novel insights into epithelial polarity proteins in Drosophila. Trends Cell Biol. 21:401–408.

Laprise, P., Beronja, S., Silva-Gagliardi, N.F., Pellikka, M., Jensen, A.M., McGlade, C.J. and Tepass, U. 2006. The FERM protein Yurt is a negative regulatory component of the Crumbs complex that controls epithelial polarity and apical membrane size. Dev Cell. 11:363–374.

Lecuit, T. and Lenne, P.F. 2007. Cell surface mechanics and the control of cell shape, tissue patterns and morphogenesis. Nat Rev Mol Cell Biol. 8:633.

Lecuit, T., and A.S. Yap. 2015. E-cadherin junctions as active mechanical integrators in tissue dynamics. Nat Cell Biol. 17:533–539.

Letunic, I., T. Doerks, and P. Bork. 2015. SMART: recent updates, new developments and status in 2015. Nucleic Acids Res. 43:D257–260.

Luo, L., Liao, Y.J., Jan, L.Y. and Jan, Y.N. 1994. Distinct morphogenetic functions of similar small GTPases: Drosophila Drac1 is involved in axonal outgrowth and myoblast fusion. Genes Dev. 8:1787–1802.

Mack, N.A. and Georgiou, M. 2014. The interdependence of the Rho GTPases and apicobasal cell polarity. Small GTPases. 5:973768.

Magie, C.R., Meyer, M.R., Gorsuch, M.S. and Parkhurst, S.M. 1999. Mutations in the Rho1 small GTPase disrupt morphogenesis and segmentation during early Drosophila development. Development. 126:5353–5364.

Manning, A.J. and Rogers, S.L. 2014. The Fog signaling pathway: insights into signaling in morphogenesis. Dev Biol. 394:6–14.

Matsuo, N., Terao, M., Nabeshima, Y.I. and Hoshino, M. 2003. Roles of STEF/Tiam1, guanine nucleotide exchange factors for Rac1, in regulation of growth cone morphology. Mol Cell Neurosci. 24:69–81.

Matsuoka, T. and Yashiro, M. 2014. Rho/ROCK signaling in motility and metastasis of gastric cancer. WJG. 20:13756.

McCormack, J., Welsh, N.J. and Braga, V.M. 2013. Cycling around cell–cell adhesion with Rho GTPase regulators. J Cell Sci. 126:379–391.

Munjal, A., J.M. Philippe, E. Munro, and T. Lecuit. 2015. A self-organized biomechanical network drives shape changes during tissue morphogenesis. Nature. 524:351–355.

Nakajima, H. and Tanoue, T. 2010. Epithelial cell shape is regulated by Lulu proteins via myosin-II. J Cell Sci. 123:555–566.

Nakajima, H. and Tanoue, T. 2011. Lulu2 regulates the circumferential actomyosin tensile system in epithelial cells through p114RhoGEF. J Cell Biol. 195:245–261.

Nakajima, H., and T. Tanoue. 2012. The circumferential actomyosin belt in epithelial cells is regulated by the Lulu2-p114RhoGEF system. Small GTPases. 3:91–96.

Ngok, S.P., Lin, W.H. and Anastasiadis, P.Z. 2014. Establishment of epithelial polarity–GEF who’s minding the GAP? J Cell Sci. 127:3205–3215.

Pédelacq, J.D., Cabantous, S., Tran, T., Terwilliger, T.C. and Waldo, G.S. 2006. Engineering and characterization of a superfolder green fluorescent protein. Nature Biotechnol. 24:79.

Pellikka, M., Tanentzapf, G., Pinto, M., Smith, C., McGlade, C.J., Ready, D.F. and Tepass, U. 2002. Crumbs, the Drosophila homologue of human CRB1/RP12, is essential for photoreceptor morphogenesis. Nature. 416:143.

Ratheesh, A. and Priya, R. 2013. Coordinating Rho and Rac: the regulation of Rho GTPase signaling and cadherin junctions. Mol Biol Transl Sci. 116:49–68.

Rauzi, M., Lenne, P.F. and Lecuit, T. 2010. Planar polarized actomyosin contractile flows control epithelial junction remodelling. Nature. 468:1110.

Ren, Y., Li, R., Zheng, Y. and Busch, H. 1998. Cloning and characterization of GEF-H1, a microtubule-associated guanine nucleotide exchange factor for Rac and Rho GTPases. J Biol Chem. 273:34954–34960.

Ridley, A.J. 2012. Historical overview of Rho GTPases. Rho GTPases. 3–12.

Rørth, P. 1998. Gal4 in the Drosophila female germline. Mech Dev. 78:113–118.

Royou, A., Sullivan, W., & Karess, R. E. 1999. In vivo studies Drosophila nonmuscle myosin II tagged with Green Fluorescent Protein. Mol Biol Cell. 10:34.

Samarin, S.N., Ivanov, A.I., Flatau, G., Parkos, C.A. and Nusrat, A. 2007. Rho/Rho-associated kinase-II signaling mediates disassembly of epithelial apical junctions. Mol Biol Cell. 18:3429–3439.

Schultz, J., Milpetz, F., Bork, P. and Ponting, C.P. 1998. SMART, a simple modular architecture research tool: identification of signaling domains. Proc Natl Acad Sci. 95:5857–5864.

Sen, A., Nagy-Zsvér-Vadas, Z. and Krahn, M.P. 2012. Drosophila PATJ supports adherens junction stability by modulating Myosin light chain activity. J Cell Biol. 199:685–698.

Shibolet, O., Giallourakis, C., Rosenberg, I., Mueller, T., Xavier, R.J. and Podolsky, D.K. 2007. AKAP13, a RhoA GTPase-specific guanine exchange factor, is a novel regulator of TLR2 signaling. J Biol Chem. 282:35308–35317.

Simões, S. D. M., Blankenship, J.T., Weitz, O., Farrell, D.L., Tamada, M., Fernandez-Gonzalez, R. and Zallen, J.A. 2010. Rho-kinase directs Bazooka/Par-3 planar polarity during Drosophila axis elongation. Dev Cell. 19:377–388.

Simões Sde, M., A. Mainieri, and J.A. Zallen. 2014. Rho GTPase and Shroom direct planar polarized actomyosin contractility during convergent extension. J Cell Biol. 204:575–589.

Strutt, D.I., Weber, U. and Mlodzik, M. 1997. The role of RhoA in tissue polarity and Frizzled signalling. Nature. 387:292.

Tanentzapf, G. and Tepass, U. 2003. Interactions between the crumbs, lethal giant larvae and bazooka pathways in epithelial polarization. Nat Cell Biol. 5:46.

Tepass, U., E. Gruszynski-DeFeo, T.A. Haag, L. Omatyar, T. Torok, and V. Hartenstein. 1996. shotgun encodes Drosophila E-cadherin and is preferentially required during cell rearrangement in the neurectoderm and other morphogenetically active epithelia. Genes Dev. 10:672–685.

Tepass, U., C. Theres, and E. Knust. 1990. crumbs encodes an EGF-like protein expressed on apical membranes of Drosophila epithelial cells and required for organization of epithelia. Cell. 61:787–799.

Tepass, U. 1996. Crumbs, a component of the apical membrane, is required for zonula adherens formation in primary epithelia of Drosophila. Dev Biol. 177:217–225.

Tepass, U. 2012. The apical polarity protein network in Drosophila epithelial cells: regulation of polarity, junctions, morphogenesis, cell growth, and survival. Annu Rev Cell Dev Biol. 28:655–685.

Tepass, U. and Knust, E. 1990. Phenotypic and developmental analysis of mutations at the crumbs locus, a gene required for the development of epithelia in Drosophila melanogaster. Dev Biol. 199:189–206.

Tepass, U. and Knust, E. 1993. Crumbs and stardust act in a genetic pathway that controls the organization of epithelia in Drosophila melanogaster. Dev Biol. 159:311–326.

Terry, S.J., Zihni, C., Elbediwy, A., Vitiello, E., San, I.V.L.C., Balda, M.S. and Matter, K. 2011. Spatially restricted activation of RhoA signalling at epithelial junctions by p114RhoGEF drives junction formation and morphogenesis. Nat Cell Biol. 13:159.

Uemura, T., H. Oda, R. Kraut, S. Hayashi, Y. Kotaoka, and M. Takeichi. 1996. Zygotic Drosophila E-cadherin expression is required for processes of dynamic epithelial cell rearrangement in the Drosophila embryo. Genes Dev. 10:659–671.

Verboon, J.M. and Parkhurst, S.M. 2015. Rho family GTPase functions in Drosophila epithelial wound repair. Small GTPases. 6:28–35.

Vichas, A., Laurie, M.T. and Zallen, J.A. 2015. The Ski2-family helicase Obelus regulates Crumbs alternative splicing and cell polarity. J Cell Biol. 211:1011–1024.

Wang, C., Shang, Y., Yu, J. and Zhang, M. 2012a. Substrate recognition mechanism of atypical protein kinase Cs revealed by the structure of PKCι in complex with a substrate peptide from Par-3. Structure. 20:791–801.

Wang, J.W., Beck, E.S. and McCabe, B.D. 2012b. A modular toolset for recombination transgenesis and neurogenetic analysis of Drosophila. PloS one. 7:42102.

Wieschaus, E., and Nüsslein-Volhard, C. (1986). In “Looking at embryos: Drosophila a practical approach” (Roberts, Ed.) pp. 199–227. IRL Press, Oxford/Washington DC.

Wodarz, A., Hinz, U., Engelbert, M. and Knust, E. 1995. Expression of crumbs confers apical character on plasma membrane domains of ectodermal epithelia of Drosophila. Cell. 82:67–76.

Zallen, J.A. and Wieschaus, E. 2004. Patterned gene expression directs bipolar planar polarity in Drosophila. Dev Cell. 6:343–355.

