## Supplemental materials for "The RhoGEF Cysts couples apical polarity proteins to Rho and myosin activity at adherens junctions"

### Supplemental Material

**Figure S1. aPKC distribution in cyst RNAi embryos.**

Staining of *cyst* RNAi embryos compared to wild-type controls for the apical marker aPKC, the junctional marker Arm, and the basolateral marker Yrt. All panels show side views of the ectoderm or epidermis at the indicated stages. Cysts in *cyst* compromised embryos are evident in stage 11, 13 and 15 embryos (arrowheads) but not at stage 10. Markers show normal subcellular distributions in cysts with apical markers facing the lumen where cuticle will be secreted (see Figure 1 C-E). Insets show triple labeled cells with aPKC shown in blue. Scale bar, 10 μm.

**Figure S2. Cyst co-aggregates with Par3 and elicits formation of ruffles and stress fibers.**

**(A)** Immunostainings of HeLa cells co-expressing various FLAG-tagged Cyst fragments and GFP::Par3. FLAG::Cyst-C and GFP::Par3 co-aggregate in cytoplasmic puncta (close-up shown in inset).

**(B)** Immunostainings of HeLa cells expressing GFP::Cyst-DH-PH with indicated markers. Dorsal and basal views are shown. Arrowheads indicate dorsal ruffles or stress fibers.

**Figure S3. Cyst interacts with Rho1.**

**(A)** Junctional levels of active Rho1 detected with PKNG58A::Venus biosensor at stage 7 and 8 in control and Cyst depleted embryos. Baz labels AJs. Notice a decrease in the levels of active Rho1 at cell-cell junctions and junctional fragmentation in Cyst depleted embryos. This Rho1 sensor also labels centrosomes (cytoplasmic bright dots), whose signal is maintained in Cyst depleted embryos as in controls and serve as internal control labeling. Scale bar, 10 μm.

**(B)** Quantification of junctional PKNG58A::Venus levels at stage 7 and 8 in control and Cyst depleted embryos. Controls: 13 embryos at stage 7 and 12 embryos at stage 8; Cyst depleted: 9 embryos at stage 7 and 4 embryos at stage 8; n=304-556 cell junctions analyzed per genotype and embryonic stage. **** $p=2.4\times10^{-207}$ and $p=1.8\times10^{-86}$.

**(C)** Cuticles are shown for the indicated genotypes. Co-expression of Cyst with DN-Rho1 partially rescues the DN-Rho1 phenotype from cuticle vesicles to a cuticle shield (dashed line) as compared to co-expression with Cyst-N, which served as a negative control. Cyst co-expression

DN-Rac1 embryo shows an additive cuticle phenotype. No modification of the Cdc42-DN phenotype is observed with Cyst co-expression.

**Figure S4. Cyst regulates actin dynamics.**

Z-projections taken from live embryos expressing Utrophin::GFP in a wild type or *cyst* shRNA background. The ventral ectoderm is shown. The loss of Cyst leads to aberrant Utrophin::GFP dynamics with an increase in Utrophin::GFP levels throughout the ectoderm. Scale bar, 10 µm. Corresponds to Movie S9 and S10.

**List of supplementary movies**

**Movie S1. Cyst is required for cell adhesion.**

Live-imaging of a wild type embryo expressing endoDEcad::GFP. Anterior is left, and ventral is down. See Figure 2 for stills and description.

**Movie S2. Cyst is required for cell adhesion.**

Live-imaging of a *cyst* RNAi embryo expressing endoDEcad::GFP. Anterior is left, and ventral is down. See Figure 2 for stills and description.

**Movie S3. AnillinRBD::GFP biosensor expression in a control embryo.**

Note accumulation of this biosensor at the adherens junctions in the ectoderm, as well as at the cytokinetic furrow and midbodies during ectodermal cell divisions, respectively. Images were acquired with a Leica SP8 resonant confocal microscope at 63× magnification, with 15-s intervals between stages 7 and 9 (3-3:50h AEL). Anterior is left, and ventral is down. See Figure 8A for stills and further description.

**Movie S4**. **AnillinRBD::GFP biosensor expression in a *cyst* RNAi embryo.**

The junctional pool of AnillinRBD::GFP is strongly reduced compared to wild-type control embryo. The probe still accumulates at the cytokinetic furrow and midbodies during and after cell division similarly to wild type. Images were acquired with a Leica SP8 resonant confocal microscope at 63× magnification, with 15-s intervals between stages 7 and 9 (3-3:50h AEL). Anterior is left, and ventral is down. See Figure 8A for stills and further description.

**Movie S5. Myosin II dynamics in a mock-RNAi control embryo.**

Embryo expressing shRNA against mCherry (mock-RNAi) and myosin II::GFP. Note two populations of myosin II, one associated with the AJs and another dynamic, pulsatile pool at the medial-apical cell cortex. Images were acquired with a Leica SP8 resonant confocal microscope at 63× magnification, with 15-s intervals between stages 7 and 10 (3-4:49h AEL). Anterior is left, and ventral is down. See Figure 9B for stills and further description.

**Movie S6. Myosin II dynamics in a *cyst* RNAi embryo.**

Embryo maternally depleted for *cyst* (mothers: *Df(cyst)*/+ *cyst* RNAi) expressing myosin II::GFP. Note a decrease in junctional and medial-apical levels of myosin II compared to control. Images were acquired with a Leica SP8 resonant confocal microscope at 63× magnification, with 15-s intervals between stages 7 and 10 (3-4:49h AEL). Anterior is left, and ventral is down. See Figure 9B for stills and further description.

**Movie S7. Myosin II dynamics in a control embryo.**

Wild-type embryo expressing Myosin II::GFP (green) and the membrane marker GAP43::mCherry (red). Note two populations of myosin II, one associated with the AJs and another dynamic, pulsatile pool at the medial-apical cell cortex. Images were acquired with a Leica SP8 resonant confocal microscope at 63× magnification, with 15-s intervals between stages 7 and 9 (3-3:58h AEL). Anterior is left, and ventral is down. See Figure 9C for stills and further description.

**Movie S8. Myosin II dynamics in a *crb* RNAi embryo.**

*crb* RNAi embryo expressing myosin II::GFP (green) and the membrane marker GAP43::mCherry (red). Note a decrease in the junctional levels of myosin II and an increase in medial-apical myosin II levels. Images were acquired with a Leica SP8 resonant confocal microscope at 63× magnification, with 15-s intervals between stages 7 and 9 (3-3:58h AEL). Anterior is left, and ventral is down. See Figure 9C for stills and further description.

**Movie S9. Cyst regulates actin dynamics.**

Live-imaging of a control embryo expressing Utrophin::GFP. The ventral ectoderm is followed from stage 10 + 1:30 hr at a time resolution of 30 sec. Movie is played at 21 fps. 63X objective. See Figure S4 for stills and description.

**Movie S10. Cyst regulates actin dynamics.**

Live-imaging of a *cyst* shRNA embryo expressing Utrophin::GFP. The ventral ectoderm is followed from stage 10 + 1:30 hr at a time resolution of 30 sec. 63X objective. Movie is played at 21 fps. See Figure S4 for stills and description.

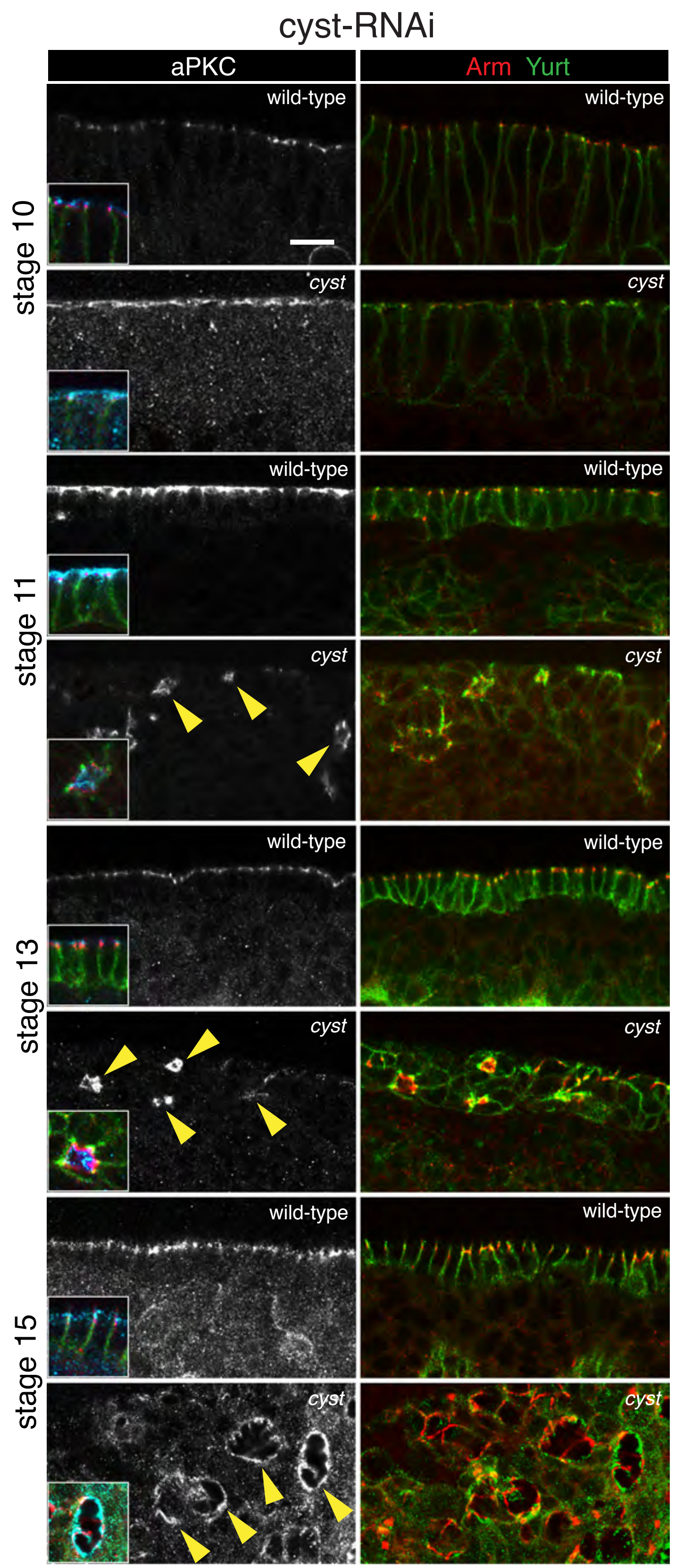


Silver et al., Figure S1

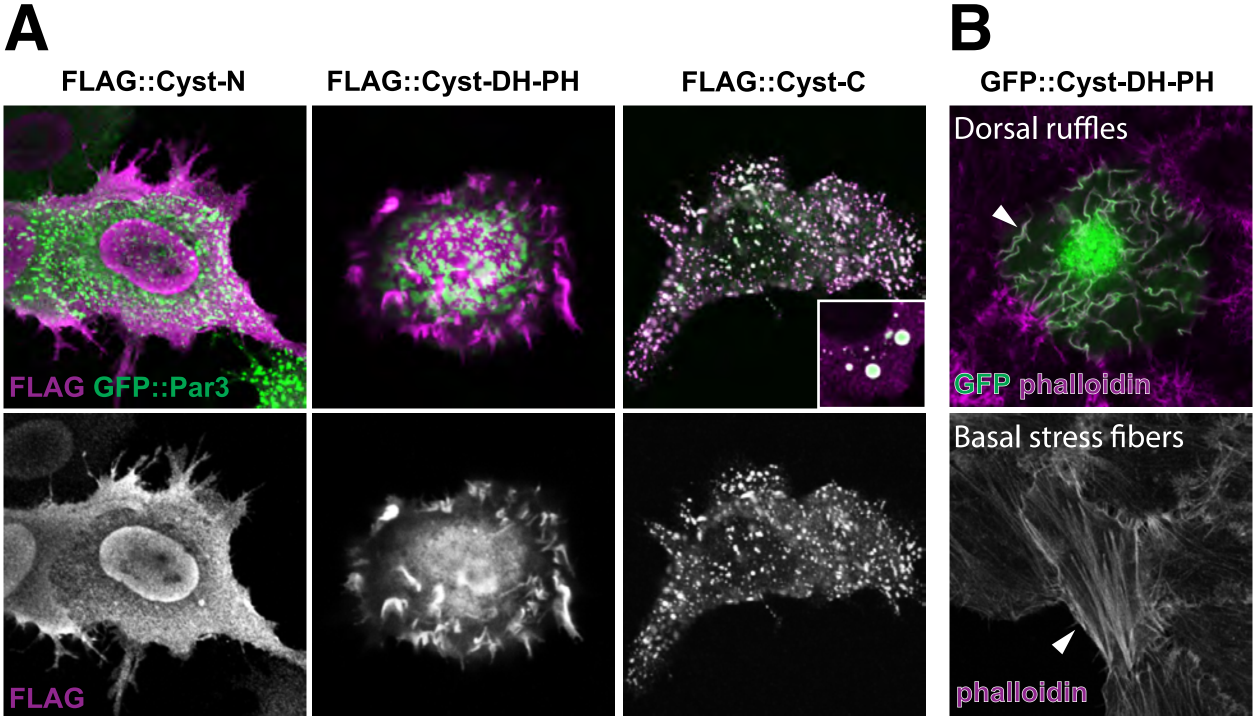


Silver et al., Figure S2

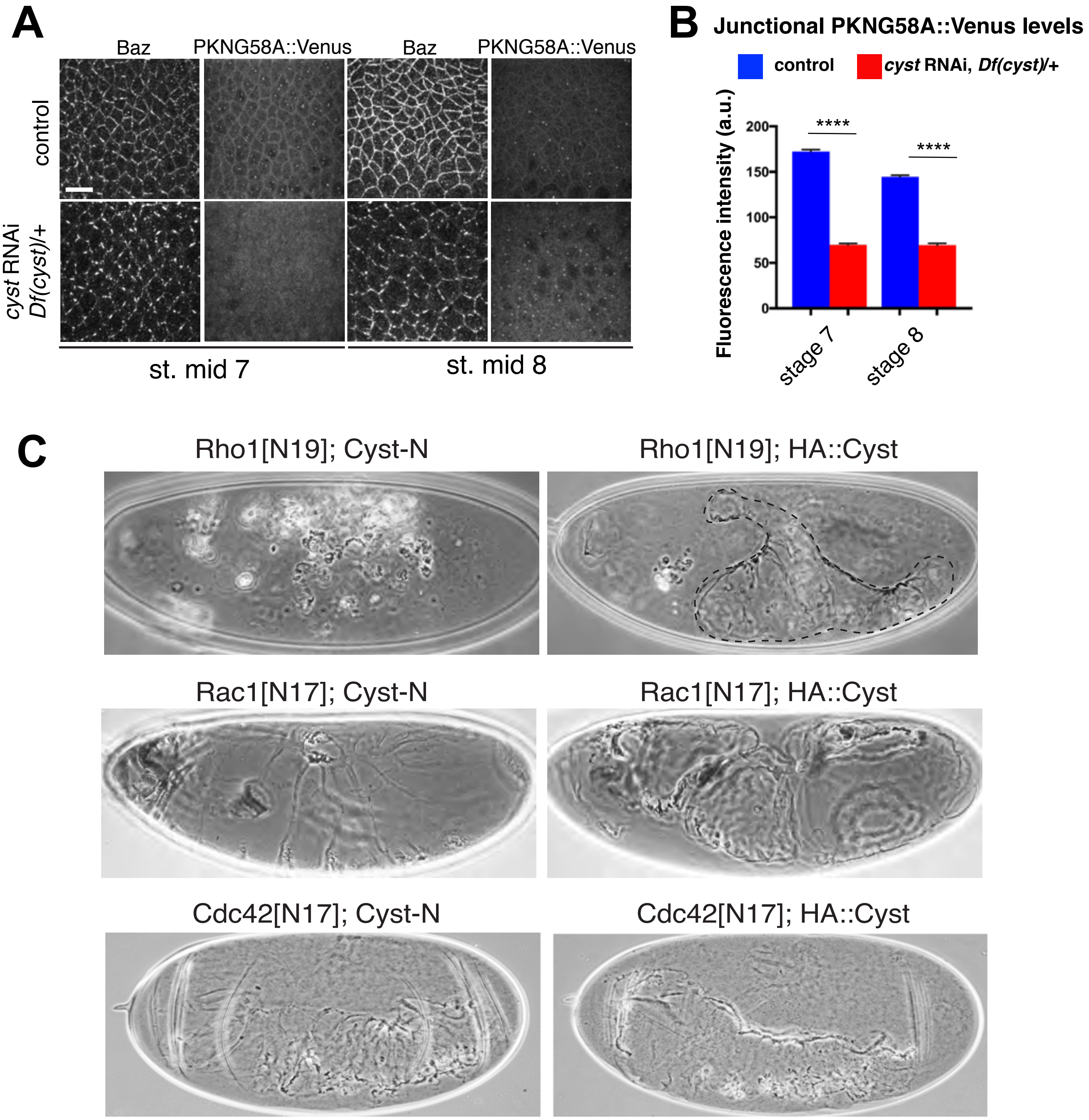


Silver et al., Figure S3

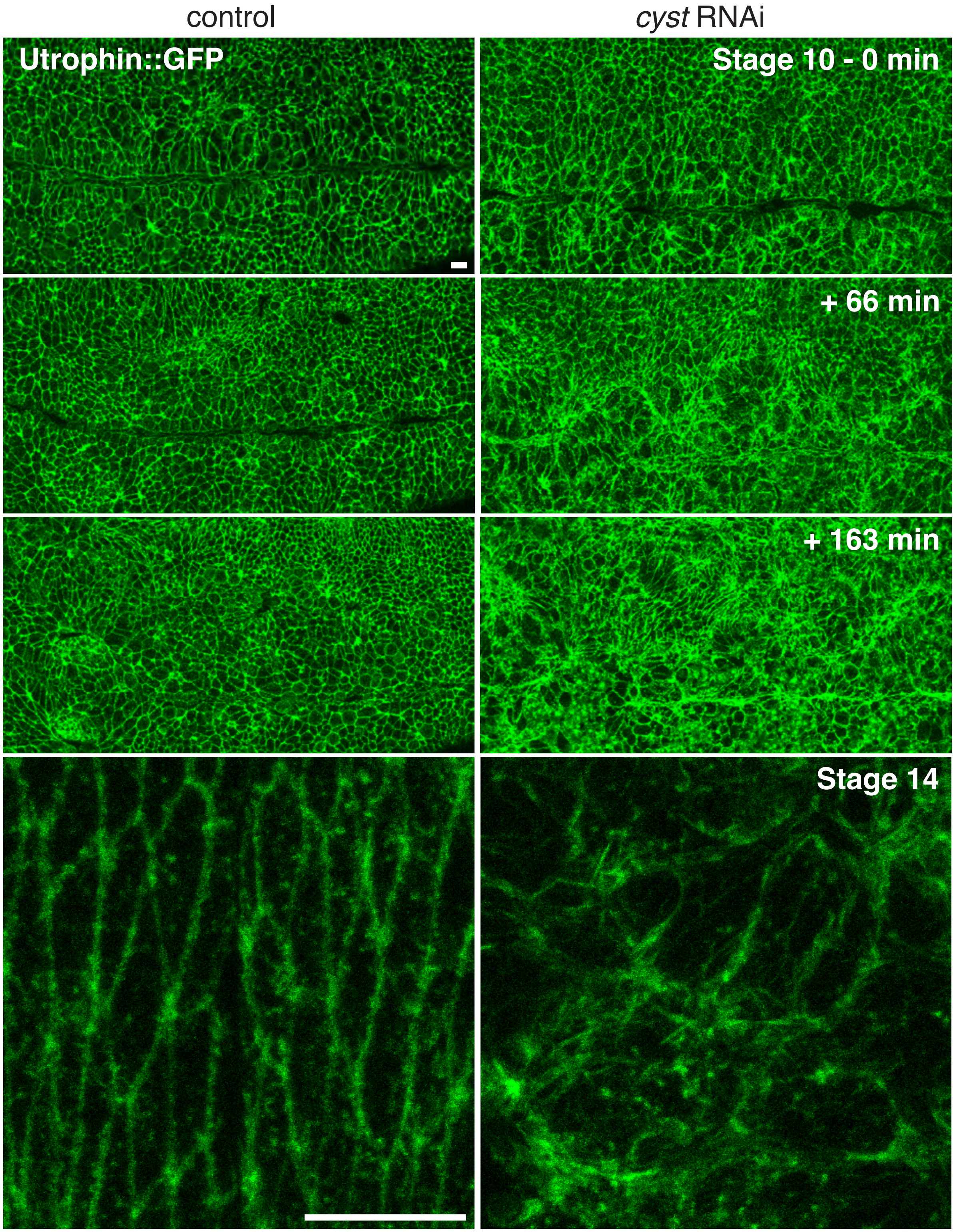


Silver et al., Figure S4
